# Positive effects of tree diversity on tropical forest restoration in a field-scale experiment

**DOI:** 10.1101/2022.09.09.507141

**Authors:** Ryan Veryard, Jinhui Wu, Michael J. O’Brien, Rosila Anthony, Sabine Both, David F.R.P. Burslem, Bin Chen, Elena Fernandez-Miranda Cagigal, H. Charles J. Godfray, Elia Godoong, Shunlin Liang, Philippe Saner, Bernhard Schmid, Yap Sau Wai, Jun Xie, Glen Reynolds, Andy Hector

## Abstract

Experiments under controlled conditions have established that ecosystem functioning is generally positively related to levels of biodiversity, but it is unclear how widespread these effects are in real-world settings and whether they can be harnessed for ecosystem restoration. We used a long-term, field-scale tropical restoration experiment to test how the diversity of planted trees affected recovery measured across a 500 ha area of selectively logged forest using multiple sources of satellite data. Replanting with species rich mixtures of tree seedlings that had higher phylogenetic and functional diversity accelerated restoration rates. Our results are consistent with a positive relationship between biodiversity and ecosystem functioning in the lowland dipterocarp rainforests of SE Asia and demonstrate that using diverse mixtures of species can enhance initial recovery after logging.

## Main Text

A quarter century of ecological experimentation has demonstrated that when other factors are held constant, ecosystem functions like biomass production are generally positively related to levels of biodiversity (*1–4*). However, for practical reasons the first generation of biodiversity manipulation experiments were conducted with systems that are relatively quick to respond, particularly communities of grassland plants (*5–8*). More recent biodiversity experiments suggest that similar diversity function relationships are present in many plantations and some forests (*9*), although there has been little research in tropical systems, particularly outside of the new world (*10–15*). It is also not clear to what degree the results of biodiversity experiments will extend to more natural settings, nor whether they can be harnessed as a nature-based solution to forest restoration and carbon capture. Here, we report early results from a field-scale experiment that tests different approaches to the restoration of lowland tropical rainforests in SE Asia, focusing in particular on the role of the diversity of tree species used for replanting. Recent results from our lowland tropical forest study system in Sabah, Malaysian Borneo, show that active restoration, including enrichment tree planting, can accelerate recovery (*16*)—here we go further in demonstrating that recovery can be enhanced by replanting with ecologically diverse mixtures of tree species.

Sabah Biodiversity Experiment (*17–19*) is designed to simultaneously test the applied question of whether increasing tree diversity in replanting schemes enhances restoration and the ecological hypothesis of whether there is a positive relationship between tree diversity and ecosystem functioning in tropical forests. There is ongoing debate over the importance of diversity for the functioning of tropical forests with some predictions of no or small ecological differences among tree species in tropical forests, and therefore an absent or weak link between diversity and functioning (*20–23*).

To be relevant to forestry and forest restoration, the Sabah Biodiversity Experiment was designed to be field-scale and covers 500 ha of selectively logged tropical forest in Malua forest reserve. The experimental treatments are applied to 4 ha plots and comprise different restoration approaches including liana removal (‘climber cutting’) and enrichment line planting where seedlings of the harvested native trees are planted into the resulting selectively logged vegetation (Fig. S1). Over 100,000 seedlings of 16 different species of the dominant dipterocarp trees (Table S1) have been planted along lines cut into the residual background vegetation left after selective logging in the 1980s and monitored periodically for survival and growth since 2002. The treatments include: unplanted controls, single-species plots enrichment planted with seedlings of one of sixteen different species of dipterocarp; polycultures planted with mixtures of 4 or 16 species; sixteen species mixtures with additional liana removal; and manipulations (within the 4-species treatment) of generic diversity and predicted canopy complexity (Table 1). To gain an overview of the effects of the experimental treatments on the whole 500 ha area of the experiment over time we used multiple sources of satellite remote sensing data including RapidEye estimates of vegetation cover, aboveground biomass and Leaf Area Index in 2012 and estimates of cover from Landsat from 1999 (prior to enrichment planting) to 2012 (*24*).

**Table 1.**
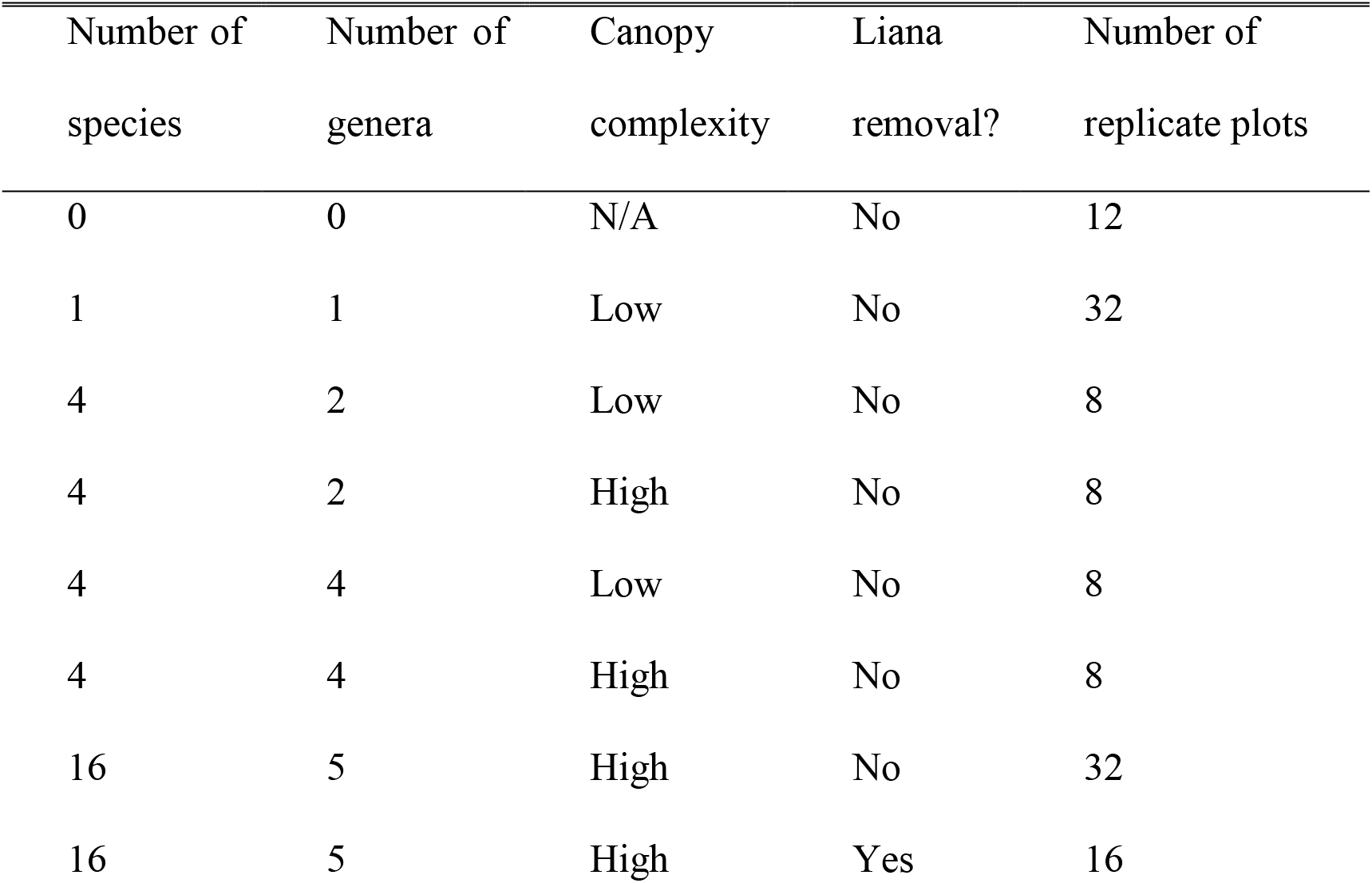
Sabah Biodiversity Experiment treatments. Treatments include number of species and genera of enrichment planted trees, predicted resulting canopy complexity, whether lianas are removed and the number of replicate plots.

Analysis of estimates of vegetation cover, aboveground biomass and Leaf Area Index derived from RapidEye satellite data in 2012 revealed several differences among the restoration treatments a decade after initial planting (Fig. 1, Table S2). Comparison of unplanted controls with enrichment planted plots revealed that active restoration increased levels of estimated biomass (Mean ± SE: 182.67 ± 4.27 vs 264.17 ± 3.883 Mg ha^-1^), cover (62.05 ± 2.28 vs 69.31 ± 2.23 %) and Leaf Area Index 4.57± 0.25 vs 5.64 ± 0.24 m^2^ m^-2^) relative to unrestored controls (Fig. 1, Table S3).

**Fig. 1.**
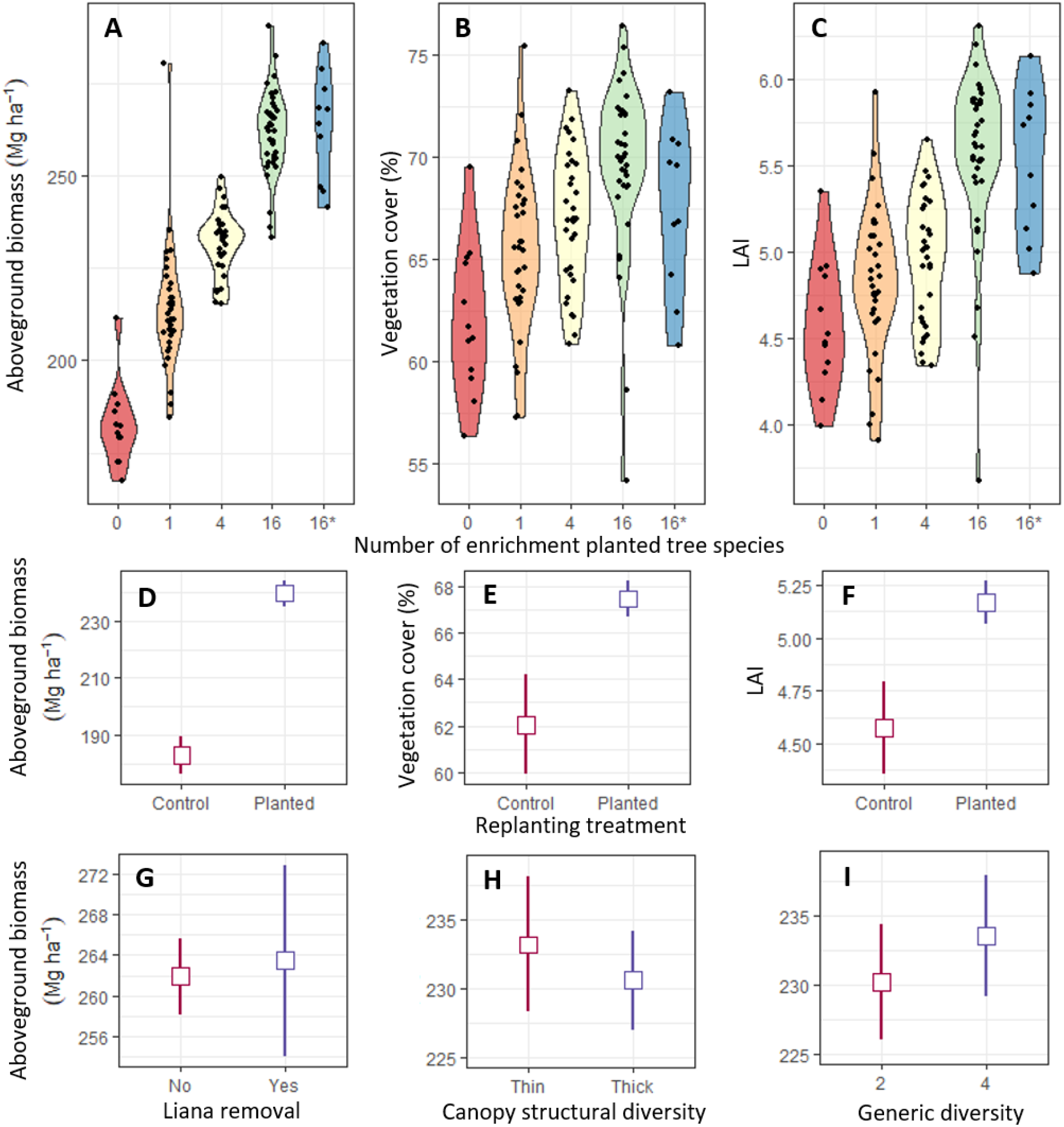
RapidEye satellite remote sensing estimates as a function of restoration treatment a decade after initial planting. (**A** to **C**) Data points for experimental plots overlaid on violin plots showing (left to right) aboveground biomass, percent vegetation cover and Leaf Area Index (LAI) in relation to enrichment planting with seedlings of 0, 1, 4, or 16 species of dipterocarp tree species (16*: enrichment planting with sixteen species plus liana cutting). (**D** to **F**) Treatment means (with 95% confidence intervals) for unplanted controls versus enrichment planted plots (panels as in top row). (**G** to **I**) Aboveground biomass as a function of (left to right) generic diversity of plots enrichment planted with four-species (2 genera vs 4 genera); canopy complexity (low vs high); and liana removal (‘climber cutting’).

While enrichment planting had a general positive effect on restoration its effectiveness was positively related to the diversity of species used. The relationship was positive and approximately linear with the logarithm of the number of enrichment-planted species: each doubling in tree species richness increased estimated biomass by 13.2 Mg ha^-1^ (± 1.5, Fig. 2, top), cover by 1.14 % (± 0.39; Fig. S2) and Leaf Area Index by 0.21 m^2^ m^-2^ (± 0.04, Fig. S3).

**Fig. 2.**
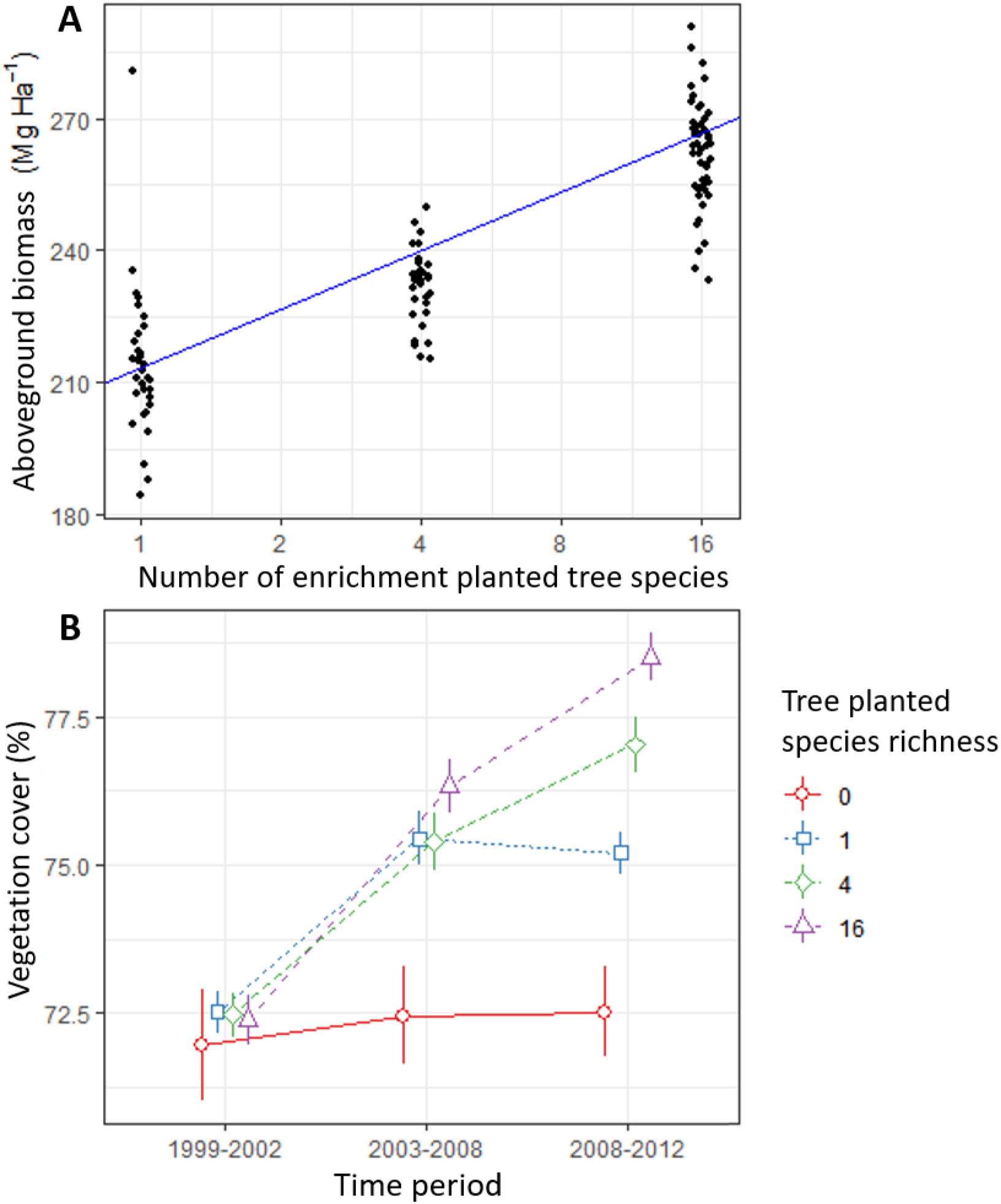
Effects of the diversity of enrichment planted trees on aboveground biomass and vegetation cover. (**A**) Estimated aboveground biomass (RapidEye) as a function of the number of enrichment-planted tree species a decade after initial planting. The line is the regression slope with the log2 number of tree species from the mixed-effects model analysis (points jittered to reduce overlap). (**B**) Changes in vegetation cover over time as a function of the number of enrichment-planted tree species. Estimates of mean cover (with 95% confidence intervals) for the LANDSAT monitoring periods 1999-2002 (prior to planting), 2003-2008 and 2008-2012 for plots enrichment planted with seedlings of 0, 1, 4, or 16 species. Individual species richness treatment levels staggered for clarity.

These treatment differences from 2012 were supported by estimates of changes in vegetation cover across three LANDSAT monitoring periods covering the preceding decade which show the absence of treatment differences prior to restoration (1999-2002), the emergence of positive effects of enrichment planting (2003-2008) and the subsequent divergence of treatments (2008-2012) with those planted with a greater diversity of tree species showing stronger recovery of vegetation cover (Fig. 2; Table S4).

Our experimental design also contains a factorial manipulation of two other aspects of diversity within the four species treatment level. Half of the four-species plots were enrichment planted with four species from four different genera and half with species from only two genera. This manipulation of generic diversity is crossed orthogonally with a treatment that compares mixtures of four species with a lower or higher diversity of predicted mature tree height that is intended to produce canopies that are thinner and simpler or thicker and more complex (Table S5). Both manipulations produced only slight increases in estimated mean aboveground biomass with enhanced generic diversity and canopy complexity (Fig. 1; Table S6) that were statistically indistinguishable between treatments (cover and leaf area index showed qualitatively similar results: Fig. S4, Table S6).

A subset of the plots planted with 16-species were also subjected to an additional treatment: reduction of lianas in the tree canopy by stem cutting (‘climber cutting’), reflecting typical Bornean forest management practice (*17*). At the time of the RapidEye data snapshot the liana removal treatment had only been applied to the southern block and the treatment had no statistically detectable effects on the satellite remote sensing estimates of biomass (Fig. 1), cover and Leaf Area Index (Fig. S4, Table S7). Previous analysis of longer-term field data (*17*) has demonstrated positive effects of liana removal on the growth and survival of trees, particularly seedlings and saplings in the understory, most likely due to increased light availability (although with potential increased seedling mortality if cutting is followed by drought). A more complete test of the liana removal treatment will require a longer series of more detailed field and remote sensing data that can discriminate between vegetation cover comprised of dipterocarp tree canopies versus lianas.

To understand why the manipulation of diversity from 1-16 species had detectible impacts on multiple measures of restoration while increasing generic diversity of the 4-species mixtures from 2 to 4 genera did not, we calculated estimates of functional and phylogenetic diversity (FD and PD) for our species mixtures (*25*). Levels of biomass were positively related to levels of FD and PD across the full species richness gradient from 1 to 16 enrichment planted species but showed only small, statistically indistinguishable increases from the two to four genera treatments and in relation to the manipulation of canopy complexity (Fig. 3). The explanation for the lack of effect of our manipulation of generic diversity probably involves both the small increase in diversity from two to four genera relative to the increase across the whole gradient from 1 to 16 species and the fact that dipterocarp taxonomy when the experiment was designed did not accurately reflect the underlying evolutionary relationships (the genus *Shorea* is now thought to be polyphyletic for example, although dipterocarp taxonomy is still unresolved).The analyses of functional diversity (FD) and phylogenetic diversity (PD) support this interpretation showing much smaller increases in diversity within the subset of treatments applied to the four species mixtures than across the entire gradient from 1 to 16 species (Fig. 3). These results suggest that the benefits of low levels of diversification in enrichment planting can be increased by the use of more species rich mixtures (at least up to the 16 species used here).

**Fig. 3.**
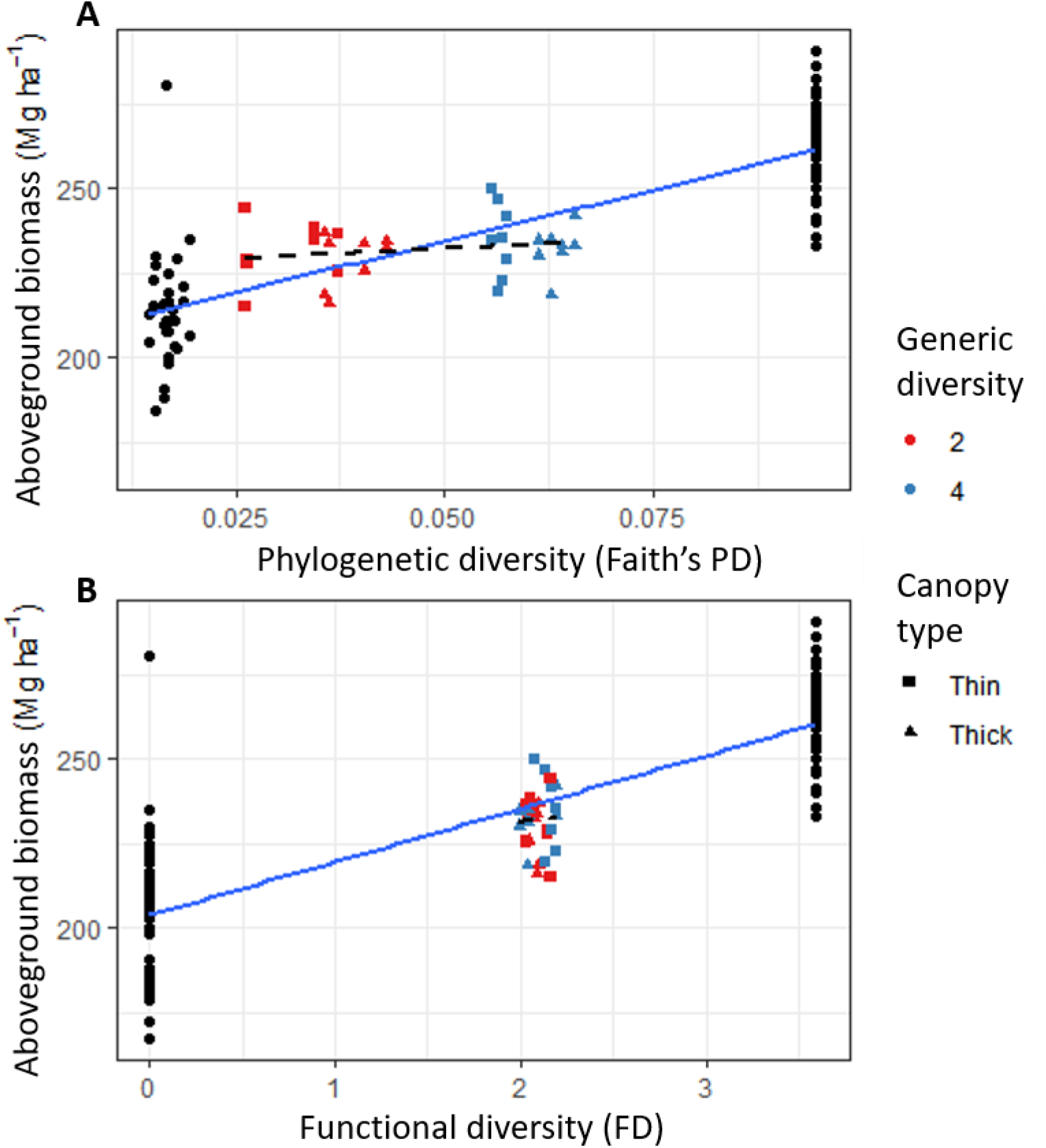
Estimated biomass as a function of phylogenetic and functional diversity. Measures of phylogenetic diversity (Faith’s PD, (**A)**) and functional diversity (FD, (**B**)) increase across the full diversity gradient from 1 to 16 species but not in relation to the treatments applied to the subset of four-species plots that manipulate generic diversity (2 vs 4 genera) and canopy complexity (lower vs higher). Solid blue lines show the positive relationship between estimated aboveground biomass and PD and FD across the full gradient from 1 to 16 species and dashed lines show the weaker, non-significant relationships for the subset of plots enrichment planted with four-species only.

Our results suggest that the positive relationship between biodiversity and ecosystem functioning observed in experiments in other ecosystems, including some forests, also applies to the lowland tropical rainforests of SE Asia. While our remote sensing data has limitations (see supplementary information: *Study limitations*) the results reported here appear robust since the same qualitative patterns are evident in two different sources of satellite data. Comparing these satellite data with field data for a similar period (*18*) suggests that during the first decade of the experiment, the effects of diversity do not come primarily through higher survival or greater trunk diameter. Instead, we hypothesize that the differences detected by satellite remote sensing are due to the development of different canopy architectures in monospecific and multi-species mixtures that we were unable to monitor in the field. Diversity-dependent growth forms have previously been shown to play a role in generating biodiversity effects in the Wageningen biodiversity experiment (*26*). Testing this hypothesis, and whether differences in canopy responses subsequently feed back to improve survival and DBH growth in mixtures, will require continued long-term monitoring and, ideally, coordinated combination of field and remote sensing data, including more detailed measurements of canopy growth.

A recent analysis of secondary succession and recovery after deforestation at sites in West Africa and Central and South America suggests forests in these areas are resilient, recovering old growth characteristic for some properties in as little as two decades (although >120 years for others) as long as land-use intensity after deforestation was low (*27*). Our results suggest the recovery of lowland forests in aseasonal SE Asia can be accelerated by active restoration through enrichment planting, especially with diverse mixtures of complementary tree species. Differences between the forests of SE Asia and other parts of the tropics are possible due to characteristics of the dominant dipterocarp species that may slow the recovery of these forests including the absence of a seedbank, intermittent mast fruiting and low dispersal ability of many species (*28–30*). Our results suggest that conservation of the diversity of tree species in these forests is needed to support the ecosystem functions and services that they provide—a matter of urgency given the recent estimate that 70% of Bornean dipterocarp species are threatened with extinction (*31*). Our results also suggest that replanting of these secondary forests with diverse mixtures of the native species removed by selective logging may provide a nature-based solution for their accelerated restoration.

## Supporting information

Data analysis supplement

## Acknowledgments

We thank all past and present research assistants for maintaining Sabah Biodiversity Experiment over the last 20 years. We acknowledge assistance and support from the South East Asia Rainforest Research partnership (SEARRP) and Sabah Forestry Department. This publication is Sabah Biodiversity Experiment article 25.

## Funding

Natural Environment Research Council grant D4T003300 (RV)

Natural Environment Research Council grant NE/K016253/1 (AH, SB, DFRPB, GR)

Comunidad de Madrid Atracción de Talento Modalidad I Fellowship 2018-T1/AMB-11095 (MJOB)

University of Zurich Research Priority Program on Global Change and Biodiversity (BS)

National Key Research and Development Program of China grant 2016YFA0600101 (JW)

## Author contributions

Conceptualization: MJOB, HCJG, AH, GR, BS, PS, RV, JW, YSW

Data curation: SB, DFRPB, BC, AH, SL, RV, JW, JX

Formal analysis: AH, RV, JW

Funding acquisition: MJOB, HCJG, AH, GR, BS, RV, JW

Investigation: SB, DFRPB, BC, AH, SL, RV, JW, JX,

Project administration: RA, MJOB, EG, AH, GR, YSW

Supervision: MJOB, EFMC, EG, AH, GR

Visualization: AH, RV, JW

Writing – original draft: AH, RV

Writing – review & editing: RA, SB, DFRPB, MJOB, BC, EFMC, HCJG, EG, AH, SL, GR, PS, BS, RV, JW, YSW, JX

## Data and materials availability

All data and code for analysis can be found at https://github.com/RVeryard/Satellite_SBE.

## List of Supplementary materials

### Supplementary materials

Materials and Methods

Supplementary Text

Tables S1 to S8

Figs. S1 to S4

References (32 – 58)

## References and Notes

1. F. Isbell, A. Gonzalez, M. Loreau, J. Cowles, S. Díaz, A. Hector, G. M. Mace, D. A. Wardle, M. I. O’Connor, J. E. Duffy, L. A. Turnbull, P. L. Thompson, A. Larigauderie, Linking the influence and dependence of people on biodiversity across scales. Nature. 546, 65–72 (2017).

2. B. J. Cardinale, J. E. Duffy, A. Gonzalez, D. U. Hooper, C. Perrings, P. Venail, A. Narwani, G. M. Mace, D. Tilman, D. A. Wardle, A. P. Kinzig, G. C. Daily, M. Loreau, J. B. Grace, A. Larigauderie, D. S. Srivastava, S. Naeem, Biodiversity loss and its impact on humanity. Nature. 486, 59–67 (2012).

3. J. S. Lefcheck, J. E. K. Byrnes, F. Isbell, L. Gamfeldt, J. N. Griffin, N. Eisenhauer, M. J. S. Hensel, A. Hector, B. J. Cardinale, J. E. Duffy, Biodiversity enhances ecosystem multifunctionality across trophic levels and habitats. Nat. Commun. 6, 1–7 (2015).

4. D. Tilman, F. Isbell, J. M. Cowles, Biodiversity and Ecosystem Functioning. Annu. Rev. Ecol. Evol. Syst. 45, 471–493 (2014).

5. A. Hector, B. Schmid, C. Beierkuhnlein, M. C. Caldeira, M. Diemer, P. G. Dimitrakopoulos, J. A. Finn, H. Freitas, P. S. Giller, J. Good, R. Harris, P. Högberg, K. Huss-Danell, J. Joshi, A. Jumpponen, C. Körner, P. W. Leadley, M. Loreau, A. Minns, C. P. H. Mulder, G. O’Donovan, S. J. Otway, J. S. Pereira, A. Prinz, D. J. Read, M. Scherer-Lorenzen, E. D. Schulze, A. S. D. Siamantziouras, E. M. Spehn, A. C. Terry, A. Y. Troumbis, F. I. Woodward, S. Yachi, J. H. Lawton, Plant diversity and productivity experiments in European grasslands. Science. 286, 1123–1127 (1999).

6. D. Tilman, P. B. Reich, J. Knops, D. Wedin, T. Mielke, C. Lehman, Diversity and productivity in a long-term grassland experiment. Science. 294, 843–845 (2001).

7. C. Roscher, V. M. Temperton, M. Scherer-Lorenzen, M. Schmitz, J. Schumacher, B. Schmid, N. Buchmann, W. W. Weisser, E. D. Schulze, Overyielding in experimental grassland communities – irrespective of species pool or spatial scale. Ecol. Lett. 8, 419–429 (2005).

8. J. van Ruijven, F. Berendse, Diversity–productivity relationships: Initial effects, long-term patterns, and underlying mechanisms. Proc. Natl. Acad. Sci. 102, 695–700 (2005).

9. Y. Feng, B. Schmid, M. Loreau, D. I. Forrester, S. Fei, J. Zhu, Z. Tang, J. Zhu, P. Hong, C. Ji, Y. Shi, H. Su, X. Xiong, J. Xiao, S. Wang, J. Fang, Multispecies forest plantations outyield monocultures across a broad range of conditions. Science. 376, 865–868 (2022).

10. D. I. Forrester, J. Bauhus, A Review of Processes Behind Diversity—Productivity Relationships in Forests. Curr. For. Reports. 2, 45–61 (2016).

11. Y. Huang, Y. Chen, N. Castro-izaguirre, M. Baruffol, M. Brezzi, A. Lang, Y. Li, X. Yang, X. Liu, K. Pei, S. Both, B. Yang, D. Eichenberg, T. Assmann, T. Behrens, X. Chen, D. Chesters, B. Ding, W. Durka, A. Erfmeier, J. Fang, M. Fischer, L. Guo, D. Guo, J. L. M. Gutknecht, J. He, C. He, A. Hector, R. Hu, A. Klein, Y. Liang, S. Li, S. Michalski, M. Scherer-lorenzen, K. Schmidt, T. Scholten, A. Schuldt, X. Shi, M. Tan, Z. Tang, S. Trogisch, Z. Wang, E. Welk, C. Wirth, T. Wubet, W. Xiang, M. Yu, X. Yu, J. Zhang, S. Zhang, N. Zhang, H. Zhou, C. Zhu, L. Zhu, H. Bruelheide, P. A. Niklaus, B. Schmid, Impacts of species richness on productivity in a large-scale subtropical forest experiment. Science. 83, 80–83 (2018).

12. J. Pelletier, A. Paquette, K. Mbindo, N. Zimba, A. Siampale, B. Chendauka, F. Siangulube, J. W. Roberts, Carbon sink despite large deforestation in African tropical dry forests (miombo woodlands). Environ. Res. Lett. 13, 094017 (2018).

13. F. Schnabel, J. A. Schwarz, A. Dănescu, A. Fichtner, C. A. Nock, J. Bauhus, C. Potvin, Drivers of productivity and its temporal stability in a tropical tree diversity experiment. Glob. Chang. Biol. 25, 4257–4272 (2019).

14. C. Messier, J. Bauhus, R. Sousa-Silva, H. Auge, L. Baeten, N. Barsoum, H. Bruelheide, B. Caldwell, J. Cavender-Bares, E. Dhiedt, N. Eisenhauer, G. Ganade, D. Gravel, J. Guillemot, J. S. Hall, A. Hector, B. Hérault, H. Jactel, J. Koricheva, H. Kreft, S. Mereu, B. Muys, C. A. Nock, A. Paquette, J. D. Parker, M. P. Perring, Q. Ponette, C. Potvin, P. B. Reich, M. Scherer-Lorenzen, F. Schnabel, K. Verheyen, M. Weih, M. Wollni, D. C. Zemp, For the sake of resilience and multifunctionality, let’s diversify planted forests! Conserv. Lett. 15, e12829 (2022).

15. E. Warner, S. C. Cook-Patton, O. T. Lewis, N. Brown, J. Koricheva, N. Eisenhauer, O. Ferlian, D. Gravel, J. S. Hall, H. Jactel, C. Mayoral, C. Meredieu, C. Messier, A. Paquette, W. C. Parker, C. Potvin, P. B. Reich, A. Hector, Higher aboveground carbon stocks in mixed-species planted forests than monocultures – a meta-analysis. bioRxiv, doi:10.1101/2022.01.17.476441.

16. C. D. Philipson, M. E. J. Cutler, P. G. Brodrick, G. P. Asner, D. S. Boyd, P. M. Costa, J. Fiddes, G. M. Foody, G. M. F. Van Der Heijden, A. Ledo, P. R. Lincoln, J. A. Margrove, R. E. Martin, S. Milne, Active restoration accelerates the carbon recovery of human-modified tropical forests. Science. 841, 838–841 (2020).

17. M. J. O’Brien, C. D. Philipson, G. Reynolds, D. Dzulkifli, J. L. Snaddon, R. Ong, A. Hector, Positive effects of liana cutting on seedlings are reduced during El Niño-induced drought. J. Appl. Ecol. 56, 891–901 (2019).

18. S. L. Tuck, M. J. O. Brien, C. D. Philipson, P. Saner, M. Tanadini, D. Dzulkifli, H. C. J. Godfray, E. Godoong, R. Nilus, R. C. Ong, B. Schmid, W. Sinun, J. L. Snaddon, M. Snoep, H. Tangki, J. Tay, P. Ulok, Y. S. Wai, M. Weilenmann, G. Reynolds, A. Hector, The value of biodiversity for the functioning of tropical forests: insurance effects during the first decade of the Sabah biodiversity experiment. Proc. R. Soc. B Biol. Sci. 283, 20161451 (2016).

19. A. Hector, C. Philipson, P. Saner, J. Chamagne, D. Dzulkifli, M. O’Brien, J. L. Snaddon, P. Ulok, M. Weilenmann, G. Reynolds, H. C. J. Godfray, The Sabah Biodiversity Experiment: a long-term test of the role of tree diversity in restoring tropical forest structure and functioning. Philos. Trans. R. Soc. B Biol. Sci. 366, 3303–3315 (2011).

20. S. P. Hubbell, A unified theory of biogeography and relative species abundance and its application to tropical rain forests and coral reefs. Coral Reefs. 16, S9–S21 (1997).

21. S. P. Hubbell, Neutral Theory and the Evolution of Ecological Equivalence. Ecology. 87, 1387–1398 (2006).

22. S. P. Hubbell, The Unified Neutral Theory of Biodiversity and Biogeography (Princeton University Press, Princeton, NJ, 2001).

23. P. B. Adler, J. HilleRislambers, J. M. Levine, A niche for neutrality. Ecol. Lett. 10, 95–104 (2007).

24. J. Wu, B. Chen, G. Reynolds, J. Xie, M. J. O’Brien, S. Liang, A. Hector, Monitoring tropical forest degradation and restoration with satellite remote sensing: A test using Sabah Biodiversity Experiment. Adv. Ecol. Res. 62, 117–146 (2020).

25. Materials and methods are available as supplementary materials.

26. J. Van Ruijven, F. Berendse, Diversity-productivity relationships: Initial effects, long-term patterns, and underlying mechanisms. Proc. Natl. Acad. Sci. U. S. A. 102, 695–700 (2005).

27. L. Poorter, D. Craven, C. C. Jakovac, M. T. van der Sande, L. Amissah, F. Bongers, R. L. Chazdon, C. E. Farrior, S. Kambach, J. A. Meave, R. Muñoz, N. Norden, N. Rüger, M. van Breugel, A. M. A. Zambrano, B. Amani, J. L. Andrade, P. H. S. Brancalion, E. N. Broadbent, H. de Foresta, D. H. Dent, G. Derroire, S. J. DeWalt, J. M. Dupuy, S. M. Durán, A. C. Fantini, B. Finegan, A. Hernández-Jaramillo, J. L. Hernández-Stefanoni, P. Hietz, A. B. Junqueira, J. K. N’dja, S. G. Letcher, M. Lohbeck, R. López-Camacho, M. Martínez-Ramos, F. P. L. Melo, F. Mora, S. C. Müller, A. E. N’Guessan, F. Oberleitner, E. Ortiz-Malavassi, E. A. Pérez-García, B. X. Pinho, D. Piotto, J. S. Powers, S. Rodríguez-Buriticá, D. M. A. Rozendaal, J. Ruíz, M. Tabarelli, H. M. Teixeira, E. V. de S. B. Sampaio, H. van der Wal, P. M. Villa, G. W. Fernandes, B. A. Santos, J. Aguilar-Cano, J. S. de Almeida-Cortez, E. Alvarez-Davila, F. Arreola-Villa, P. Balvanera, J. M. Becknell, G. A. L. Cabral, C. Castellanos-Castro, B. H. J. de Jong, J. E. Nieto, M. M. Espírito-Santo, M. C. Fandino, H. García, D. García-Villalobos, J. S. Hall, A. Idárraga, J. Jiménez-Montoya, D. Kennard, E. Marín-Spiotta, R. Mesquita, Y. R. F. Nunes, S. Ochoa-Gaona, M. Peña-Claros, N. Pérez-Cárdenas, J. Rodríguez-Velázquez, L. S. Villanueva, N. B. Schwartz, M. K. Steininger, M. D. M. Veloso, H. F. M. Vester, I. C. G. Vieira, G. B. Williamson, K. Zanini, B. Hérault, Multidimensional tropical forest recovery. Science. 374, 1370–1376 (2021).

28. J. Ghazoul, Dipterocarp biology, ecology, and conservation (Oxford University Press, 2016).

29. L. F. Banin, E. H. Raine, L. M. Rowland, R. L. Chazdon, S. W. Smith, N. E. B. Rahman, Butler, C. Philipson, G. G. Applegate, P. Axelsson, S. S. Budiharta, S. C. Chua, M. E. Cutler, S. Elliott, E. Gemita, E. Godoong, L. L. Graham, R. M. Hayward, A. A. Hector, U. Ilstedt, J. Jensen, S. Kasinathan, C. J. Kettle, D. Lussetti, B. Manohan, C. Maycock, K. M. Ngo, M. J. O’Brien, A. Osuri, G. Reynolds, Y. Sauwai, S. Scheu, M. Silalahi, E. M. Slade, T. Swinfield, D. A. Wardle, C. Wheeler, K. L. Yeong, D. F. Burslem, Philos. Trans. R. Soc. B Biol. Sci., in press.

30. L. Banin, S. L. Lewis, G. Lopez-Gonzalez, T. R. Baker, C. A. Quesada, K. J. Chao, D. F. R. P. Burslem, R. Nilus, K. Abu Salim, H. C. Keeling, S. Tan, S. J. Davies, A. Monteagudo Mendoza, R. Vásquez, J. Lloyd, D. A. Neill, N. Pitman, O. L. Phillips, Tropical forest wood production: a cross-continental comparison. J. Ecol. 102, 1025–1037 (2014).

31. Botanic Gardens Conservation International, “State of the World’s Trees” (Botanic Gardens Conservation International, Richmond, UK, 2021).

32. D. Lussetti, E. P. Axelsson, U. Ilstedt, J. Falck, A. Karlsson, Supervised logging and climber cutting improves stand development: 18 years of post-logging data in a tropical rain forest in Borneo. For. Ecol. Manage. 381, 335–346 (2016).

33. C. W. Marsh, A. G. Greer, Forest land-use in Sabah, Malaysia: an introduction to Danum Valley. Philos. Trans. - R. Soc. London, B. 335, 331–339 (1992).

34. P. Saner, thesis, University of Zurich, Zurich, Switzerland (2009).

35. T. C. Whitmore, Tropical rain forests of the far east (Clarendon Press, Oxford, ed. 2nd, 1984).

36. F. Q. Brearley, L. F. Banin, P. Saner, The ecology of the Asian dipterocarps. Plant Ecol. Divers. 9, 429–436 (2016).

37. S. Numata, M. Yasuda, R. O. Suzuki, T. Hosaka, N. S. M. Noor, C. D. Fletcher, M. Hashim, Geographical Pattern and Environmental Correlates of Regional-Scale General Flowering in Peninsular Malaysia. PLoS One. 8, e79095 (2013).

38. Y. Hautier, P. Saner, C. Philipson, R. Bagchi, R. C. Ong, A. Hector, Effects of Seed Predators of Different Body Size on Seed Mortality in Bornean Logged Forest. PLoS One. 5, e11651 (2010).

39. M. J. O’Brien, D. F. R. P. Burslem, A. Caduff, J. Tay, A. Hector, Contrasting nonstructural carbohydrate dynamics of tropical tree seedlings under water deficit and variability. New Phytol. 205, 1083–1094 (2015).

40. P. S. Ashton, in Flora Malesiana (1982), pp. 237–552.

41. P. Saner, Y. Y. Loh, R. C. Ong, A. Hector, Carbon Stocks and Fluxes in Tropical Lowland Dipterocarp Rainforests in Sabah, Malaysian Borneo. PLoS One. 7, e29642 (2012).

42. J. O. Sexton, X. P. Song, M. Feng, P. Noojipady, A. Anand, C. Huang, D. H. Kim, K. M. Collins, S. Channan, C. DiMiceli, J. R. Townshend, Global, 30-m resolution continuous fields of tree cover: Landsat-based rescaling of MODIS vegetation continuous fields with lidar-based estimates of error. Int. J. Digit. Earth. 6, 427–448 (2013).

43. P. W. Miller, A. Kumar, T. L. Mote, F. D. S. Moraes, D. R. Mishra, Persistent Hydrological Consequences of Hurricane Maria in Puerto Rico. Geophys. Res. Lett. 46, 1413–1422 (2019).

44. C. Li, H. Wulf, B. Schmid, J. S. He, M. E. Schaepman, Estimating plant traits of alpine grasslands on the qinghai-tibetan plateau using remote sensing. IEEE J. Sel. Top. Appl. Earth Obs. Remote Sens. 11, 2263–2275 (2018).

45. B. Brede, J. P. Gastellu-Etchegorry, N. Lauret, F. Baret, J. G. P. W. Clevers, J. Verbesselt, M. Herold, Monitoring Forest Phenology and Leaf Area Index with the Autonomous, Low-Cost Transmittance Sensor PASTiS-57. Remote Sens. 10, 1032 (2018).

46. G. Tyc, J. Tulip, D. Schulten, M. Krischke, M. Oxfort, The RapidEye mission design. Acta Astronaut. 56, 213–219 (2005).

47. F. Jung-Rothenhäusler, H. Weichelt, M. Pach, RapidEye – A Novel Approach to Space Borne Geo-Information Solutions. ISPRS Hann. Work. 2007, 4–7 (2007).

48. G. Chander, M. O. Haque, A. Sampath, A. Brunn, G. Trosset, D. Hoffmann, S. Roloff, M. Thiele, C. Anderson, Radiometric and geometric assessment of data from the RapidEye constellation of satellites. Int. J. Remote Sens. 34, 5905–5925 (2013).

49. M. Pfeifer, L. Kor, R. Nilus, E. Turner, J. Cusack, I. Lysenko, M. Khoo, V. K. Chey, A. C. Chung, R. M. Ewers, Mapping the structure of Borneo’s tropical forests across a degradation gradient. Remote Sens. Environ. 176, 84–97 (2016).

50. J. Qi, A. Chehbouni, A. R. Huete, Y. H. Kerr, S. Sorooshian, A modified soil adjusted vegetation index. Remote Sens. Environ. 48, 119–126 (1994).

51. J. A. Gallardo-Cruz, J. A. Meave, E. J. González, E. E. Lebrija-Trejos, M. A. Romero-Romero, E. A. Pérez-García, R. Gallardo-Cruz, J. L. Hernández-Stefanoni, C. Martorell, Predicting Tropical Dry Forest Successional Attributes from Space: Is the Key Hidden in Image Texture? PLoS One. 7, e30506 (2012).

52. M. Vellend, W. K. Cornwell, K. Magnuson-Ford, A. ø. Mooers, in Biological Diversity: Frontiers in Measurement and Assessment, A. Magurran, B. McGill, Eds. (Oxford Univ. Press, 2010), pp. 194–207.

53. O. L. Petchey, K. J. Gaston, Functional diversity (FD), species richness and community composition. Ecol. Lett. 5, 402–411 (2002).

54. F. Barragán, C. E. Moreno, F. Escobar, G. Halffter, D. Navarrete, Negative Impacts of Human Land Use on Dung Beetle Functional Diversity. PLoS One. 6, e17976 (2011).

55. E. Laliberté, P. Legendre, A distance-based framework for measuring functional diversity from multiple traits. Ecology. 91, 299–305 (2010).

56. C. C. P. Cosset, D. P. Edwards, The effects of restoring logged tropical forests on avian phylogenetic and functional diversity. Ecol. Appl. 27, 1932–1945 (2017).

57. B. F. Chabot, D. J. Hicks, The ecology of leaf life spans. Annu. Rev. Ecol. Syst. 13, 229–259 (1982).

58. S. J. Wright, K. Kitajima, N. J. B. Kraft, P. B. Reich, I. J. Wright, D. E. Bunker, R. Condit, J. W. Dalling, S. J. Davies, S. Díaz, B. M. J. Engelbrecht, K. E. Harms, S. P. Hubbell, C. O. Marks, M. C. Ruiz-Jaen, C. M. Salvador, A. E. Zanne, Functional traits and the growth — mortality trade-off in tropical trees. Ecology. 91, 3664–3674 (2010).

