## Supplementary methods for "Positive effects of tree diversity on tropical forest restoration in a field-scale experiment"

**This PDF file includes:**

Materials and Methods
Supplementary Text
Figs. S1 to S4
Tables S1 to S8

### Materials and Methods:

Study system

The Sabah Biodiversity Experiment (http://www.sabahbiodiversityexperiment.org) occupies 500 ha in the southern part of the Malua Forest Reserve, in Sabah, Malaysian Borneo (Fig. S1). The Malua Forest Reserve is an area of approximately 35,000 ha of predominantly selectively logged forest that is publicly owned through Yayasan Sabah (The Sabah Foundation), which holds a 100-year concession under its goal to increase socioeconomic standards in the state. Within the wider Yayasan Sabah logging concession is the Innoprise-FACE Foundation Rainforest Rehabilitation project (INFAPRO), a 25,000 ha area dedicated to promoting the rehabilitation of forests through large scale enrichment planting within logged areas. To help provide practical recommendations, the Sabah Biodiversity Experiment followed INFAPRO enrichment planting techniques. The region experiences an average temperature of 27 $^{o}$C and an annual rainfall of >3000 mm, distributed between two wet seasons (*32–34*). The Malua Forest Reserve area has been logged twice, once in the 1980s and again in 2007. The 500 ha area of the Sabah Biodiversity Experiment itself was spared the second round of selective logging in 2007 due to the establishment of the experiment in 2002 and has therefore been recovering from the initial round of logging for nearly 40 years. Elevation at this site is under 250 m, with 0-20$^{o}$ range in topography. The pre-logging timber volume of this region has been estimated at 193-221 $m^3$ $ha^{-1}$, of which dipterocarps account for the vast majority at 180-216 $m^3$ $ha^{-1}$ (*19*).

Study species

The 16 species used in this experiment are native species belonging to the Dipterocarpaceae: *Dipterocarpus conformis* Slooten, *Dryobalanops lanceolata* Burck, *Hopea ferruginea* Parij, *Hopea sangal* Korth., *Parashorea malaanonan* (Blanco) Merr., *Parashorea tomentella* (Blanco) Merr., *Shorea argentifolia* Sym., *Shorea beccariana* Bruck, *Shorea faguetiana* Heim., *Shorea gibbosa* Brandis., *Shorea johorensis* Foxw., *Shorea leprosula* Miq., *Shorea macrophylla* Ashton, *Shorea macroptera* King, *Shorea ovalis* Korth., and *Shorea parvifolia* Dyer. Generally, dipterocarps in Sabah are emergent tree species, rarely found 1,200 m above sea level (*35*). They have an array of characteristic which likely contribute to their dominance in SE Asian forests including their symbiotic ectomycorrhizal associations (*36*), and wind-dispersed winged fruits (*37*). Reproduction takes place largely through 'mast fruiting' events that occur between 2-10 years apart, where many or most of the dipterocarp species simultaneously produce fruit. Dipterocarps have recalcitrant seeds (*38*) and no soil seed bank (*16*). Instead, successful recruits form a seedling bank which often suffer from heavy herbivory (*39*). Dipterocarps dominate the lowland forests of SE Asia in terms of biomass but have been heavily selectively logged (*40*).

Experimental design

Sabah Biodiversity Experiment features several experimental treatments within its replicated, randomised block design. The experiment consists of 124 four-hectare (200 m x 200 m) plots, divided into two blocks separated by an old logging road (60 plots in the north block and 64 to the south). Each plot (apart from the unplanted controls) is enrichment planted with a mixture of seedlings with a controlled species number (richness) and composition. The design ensures at least one replicate plot for each species richness and composition treatment level in each of the two blocks. Each plot contains 20 parallel planting lines, separated by 10 m areas of remnant vegetation left after the prior selective logging. Within each line, seedlings were planted with 3 m spacing, and planting lines were initially cleared of bamboo, lianas, and shrubs up to a maximum of 1 m either side of the line of planted seedlings. The experiment was primarily designed to manipulate the diversity and composition of enrichment planted dipterocarps, but also investigates the forest management practice of liana removal ('climber cutting'). 114 of the plots make up a gradient in the diversity of enrichment-planted tree species comprising mixtures of 1, 4, or 16 species. The remaining 12 plots were left as naturally regenerating unplanted controls (six in each block). The design uses a set of 16 species that were available in the local seedling nursery in sufficient numbers. These 16 species were grown in single-species enrichment planting 'monocultures' and combined together to form enrichment planting 'polycultures' of 4 or 16 species (Table 1). The plots enrichment planted with only a single species of dipterocarp allow a comparison of individual species identity effects since each species has a replicate in each of the two blocks (a total of 32 1-species plots).

The intermediate 4-species diversity level is comprised of 16 different species compositions that produce two further treatments that are factorially crossed. These two treatments manipulate generic diversity (two levels) and predicted canopy structural diversity (two levels). The generic diversity treatment compares mixtures of four species comprising two or four dipterocarp genera. The canopy structural diversity treatment also features two levels that either combine species with similar predicted mature heights or with a wider range of these predicted values. In total, this factorial manipulation of generic and structural diversity comprises 32 plots (the $2^2$ factorial combination of the 4 treatments, each with 4 replicate species compositions, each replicated in the two blocks) (Table S5).

Sixteen plots of the most diverse (16-species) mixtures underwent two rounds of liana removal (‘climber cutting’), which were compared with 32 plots enrichment-planted with the same number of species but without this local climber cutting restoration strategy (*17*). Due to practical constraints these cuttings took place in two stages. In July 2011 ten plots were cut in the southern block, and in June 2014 these ten plots, as well as six plots in the northern block, underwent a full round of liana removal. Therefore, at the time of the RapidEye satellite remote sensing in 2012 only the ten plots in the southern block had been subjected to the liana removal treatment. Nevertheless, to avoid the risk of missing effects of this treatment, we included it in the statistical analysis.

In line with standard enrichment planting procedure, after the initial cohort of seedlings were planted (between January 2002 and September 2003), a second cohort was planted to replace initial mortalities (cohort 2 planted September 2008 to August 2009). In combination, the two cohorts planted and surveyed a total of 96,369 dipterocarp seedlings. Further details of the Malua reserve and Sabah Biodiversity Experiment can be found in previous publications (*18*, *19*, *41*).

<u>Remote sensing</u>
Landsat Vegetation Continuous Fields (VCF) tree cover, RapidEye and MODIS imagery were selected to estimate variation in canopy structure based on the needs of data accuracy, the size of the study site and plots and the time period of the experiment (Table S8).

<u>Landsat Vegetation Continuous Fields tree cover</u>

The Landsat Continuous Fields tree cover (Landsat tree cover) estimates the percentage of horizontal ground per 30 m pixel which is covered with vegetation of at least 5 m vertical height (*42*). In this study we refer to Landsat tree cover as Landsat vegetation cover, as in Sabah virtually all vegetation detected by Landsat is higher than this minimum. The product is derived from all 7 bands of Landsat-5 Thematic Mapper (TM) and/or Landsat Enhanced Thematic Mapper Plus (ETM+). The partial resolution of the Landsat vegetation cover dataset is 30 m, which is appropriate for the Sabah Biodiversity Experiment's plot size of 200 m x 200 m, giving c. 44 pixels per plot. This dataset contains three epochs, 2000, 2005, and 2010, each consisting of a composite of several years' worth of images in order to minimise the effects of cloud cover. The 2000 epoch consists of data from 1999 to 2002, our 2005 epoch contains years 2003 to 2008, and the 2010 epoch ranges 2008 to 2012.

<u>MODIS MCD15A3H</u>
MODIS MCD15A3H Leaf Area Index (LAI) is widely used in forest monitoring and exhibits very high accuracy (*43–45*). However, the spatial resolution of 500 m means that each plot does not even have a single complete pixel. Instead, a comparison of the entire SBE sites with the surrounding re-logged area is reported elsewhere (*24*).

<u>RapidEye imagery</u>
This study used a RapidEye satellite image of the Sabah Biodiversity Experiment site for August 2012. RapidEye imagery uses a higher-spatial resolution of 5 m and a temporal resolution of 5.5 days (*46*, *47*). This multi-spectral scanner of the RapidEye satellites acquires data in five bands. The blue (0.44-0.51 µm), green (0.52-0.59 µm), red (0.63-0.68 µm), and near-infrared (0.76-0.85 µm) are very similar to that of the Landsat Spectral band equivalents, while also having an additional red-edge band (0.69-0.73 µm). This band allows RapidEye satellite images to provide greater sensitivity to spatiotemporal changes in vegetation (*48*, *49*).

<u>Vegetation metrics inversion from RapidEye image</u>
A FLAASH atmospheric correction model was applied to the RapidEye image, and vegetation cover, LAI, and aboveground biomass (AGB) were calculated using empirical formulae develop in Pfeifer et al. (*49*) (Eqs. S1, S2, and S3, respectively), which used RapidEye imagery of the nearby SAFE landscape. Although these equations were not derived for Malua (where SBE is located), the SAFE landscape is close by, and significantly more so than all other options. This provided us with high resolution (5 m for RapidEye) estimates of (LAI), vegetation cover and AGB using a method developed and validated for lowland dipterocarp forests in the same part of Sabah. Further details of the inversion methodology used can be found in the previous publication (*49*).

**Eqn. S1.**

$$\text{Vegetation cover} = 2.66 - 0.66 \cdot \text{Red} + 0.3 \cdot \text{RedEdge} - 0.08 \cdot \text{NearIR} - 0.17 \cdot \text{DissB3} + 1.48 \cdot \text{DissB4} - 0.42 \cdot \text{DissB5}$$

**Eqn. S2.**

$$LAI = 0.9 - 0.59 \cdot Red + 0.41 \cdot RedEdge - 0.11 \cdot NearIR - 0.53 \cdot DissB3 + 1.08 \cdot DissB4 - 0.36 \cdot DissB5$$

**Eqn. S3.**

$$AGB = 19.45 - \exp(MSAV12) - 2.39 \cdot Green + 1.08 \cdot RedEdge + 2.65 \cdot DissB2 - 0.28 \cdot DissB3 + 0.09 \cdot DissB4 - 0.13 \cdot DissB5$$

Where MSAV12 is the Modified Soil-Adjusted Vegetation Index 2 (*50*). Green, Red, RedEdge, and NearIR all correspond to the RapidEye bands of the same name, and DissB2, DissB3, DissB4, and DissB5 are the grey-level dissimilarities of green band, red band, near-infrared band, and red-edge band, respectively (*51*). Satellite imagery were pre-processed using ArcGIS and overlaid with the SBE plot layout based on GPS coordinates collected for the perimeter of each block.

Phylogenetic and functional diversity

To further investigate the effect of genetic diversity within the four-species mixtures on estimated index values, we calculated Faith's phenotypic diversity (PD) (*52*), defined as the total branch length of the minimum spanning tree from each node to the tree root. We also calculated functional diversity (FD), a functional equivalent of Faith's PD, by the sum of branch lengths on a functional dendrogram (*53*). Out of 46 total trait measurements available we opted *a priori* to use leaf nitrogen, leaf phosphorus, leaf thickness, dry weight, wood density, and specific leaf area as these measurements have been used to estimate FD in the literature most widely (*54–56*), and there are current associations with the trade-off between rapid resource acquisition and faster growth and enhanced environmental tolerance and reduced growth (*57*, *58*). This also avoided using a large number of partially correlated variables. Both PD and FD measurements are dependent on species richness, and so contain information of phylogenetic and functional diversity both between and within species richness levels.

Statistical analysis

We analysed the satellite remote sensing data using linear mixed-effects models that implemented a series of *a priori* contrasts investigating the effects of the following experimental treatments:

1) Replanting (unplanted versus enrichment planted plots)
2) Species richness of enrichment planted trees (1, 4, or 16 species)
3) Generic richness (1, 2, 4, or 5 genera across the whole gradient and 2 vs 4 genera within the 4-species plots)
4) Predicted canopy complexity (mixtures with similar adult heights vs a diversity of adult tree heights)
5) Liana removal ('climber cutting'; 16-species plots with and without lianas removed)

The linear mixed-effects models were implemented using the lme4 package (version 1.1-26) for R (version 4.0.5). As would be expected, the RapidEye estimates of vegetation cover, LAI, and AGB values were positively correlated, and so to reduce the number of statistical tests, we focused on AGB as that is the most widely used variable in the current relevant literature and the variable most often communicated to, and understood by, managers and other relevant stakeholders. For our primary questions we fitted two linear mixed effects models. All contained RapidEye Biomass estimation as the primary response variable.

**Model 1a**

```
y ~ planting + species diversity + generic diversity + canopy complexity + liana removal + (1|Block) + (1|Spp_comp)
```

Where y is a continuous response variable (here aboveground biomass, vegetation cover or LAI), planting is a fixed factor with two levels (unplanted vs planted), species diversity is a fixed factor with 4 levels: 0 (unplanted controls), 1, 4, or 16 species, generic diversity is a fixed factor with 2 levels (2 or 4 genera), canopy complexity is a fixed factor with two levels (high or low diversity of predicted adult tree height), liana removal is a fixed factor with two levels (lianas removed or not removed), Block is a random factor with 2 levels (northern block or southern block), Species composition is a random factor with 33 levels, and (1|Factor) indicates random factors specified to have random intercepts. Note that the leading contrast in this model for unplanted vs planted plots already accounts for the comparison of species with 0 vs 1-16 enrichment planted species.

**Model 1b**

```
y ~ planting + log2(species richness) + species diversity + generic diversity + canopy complexity + liana removal + (1|Block) + (1|Spp_comp)
```

Where all terms are as defined in model 1a except $\log_2$(species richness), which is a continuous fixed response variable for the ($\log_2$-transformed) number of enrichment planted tree species. This variable captures planted species richness from 1-16 species and does not include unplanted controls, which are captured in the planting component of model 1b. Species diversity has been fitted sequentially after species richness to capture deviations from log-linearity.

Despite its various advantages the RapidEye data is for a single period in August 2012. To confirm these differences and investigate their development over time we used the three epochs of Landsat imagery covering the total period 1999-2012, extending the mixed-effects model for the RapidEye data to incorporate the effects of the three time epochs present in the Landsat data:

**Model 2**

```
y ~ planting + species richness * year + generic diversity +
canopy complexity + treatment + (1|Block) + (1|Spp_comp) +
(1|Plot)
```

Where all terms are as defined in models 1a and 1b except year, which is a fixed factor with 3 levels and where * indicates an interaction between two variables. Since there are three repeated measures per plot, one for each epoch, we also added a random factor for Plot (with 124 levels). To analyse the influence of functional or phylogenetic diversity we fitted copies of model 1a and 1b, but changing Generic diversity for either PD or FD.

**Supplementary Text**

Study Limitations

<u>Liana removal and RapidEye imagery data comparison</u>

As of August 2012, only the southern block had undergone a round of liana removal. As such the six northern block plots are *de facto* untreated 16-species plots for this purposes of this study, and were classed as such when it came to relevant analyses.

<u>Landsat and RapidEye cover comparison</u>

One immediate comparison is that Landsat cover estimates are greater than RapidEye cover estimates, by a consistent amount (9.76 ± 0.371 %), irrespective of plot treatment, with a Pearson product-moment correlation of 0.389 ($p = 7x10^{-6}$, df = 122). There are three non-mutually exclusive explanations for this difference. Firstly, the spatial resolution of Landsat vegetation cover is much lower than that of RapidEye which in theory means that more information can be extracted. However, it may be more difficult to extract accurately from high spatial resolution data because more kinds of information can appear with an increase in spatial resolution which would otherwise be neglected or offset with a lower spatial resolution, such as topography. Secondly, Landsat vegetation cover is a remote sensing production. Values from the Landsat 2010 epoch were collected from a range of years 2008-2012, whilst the RapidEye image was collected from a single time point in 2012. This would certainly lead to some variation in vegetation cover between both images. Thirdly, the calculation method of Landsat VC is developed for global-coverage, while that of RapidEye VC is developed specifically for Borneo. Theoretically the latter is more credible, but has far fewer users and no information from SBE to be used to validate. Overall, we cannot determine which index performs better, and so to be as consistent as possible Landsat imagery was used in this analysis only when multiple time points were required.

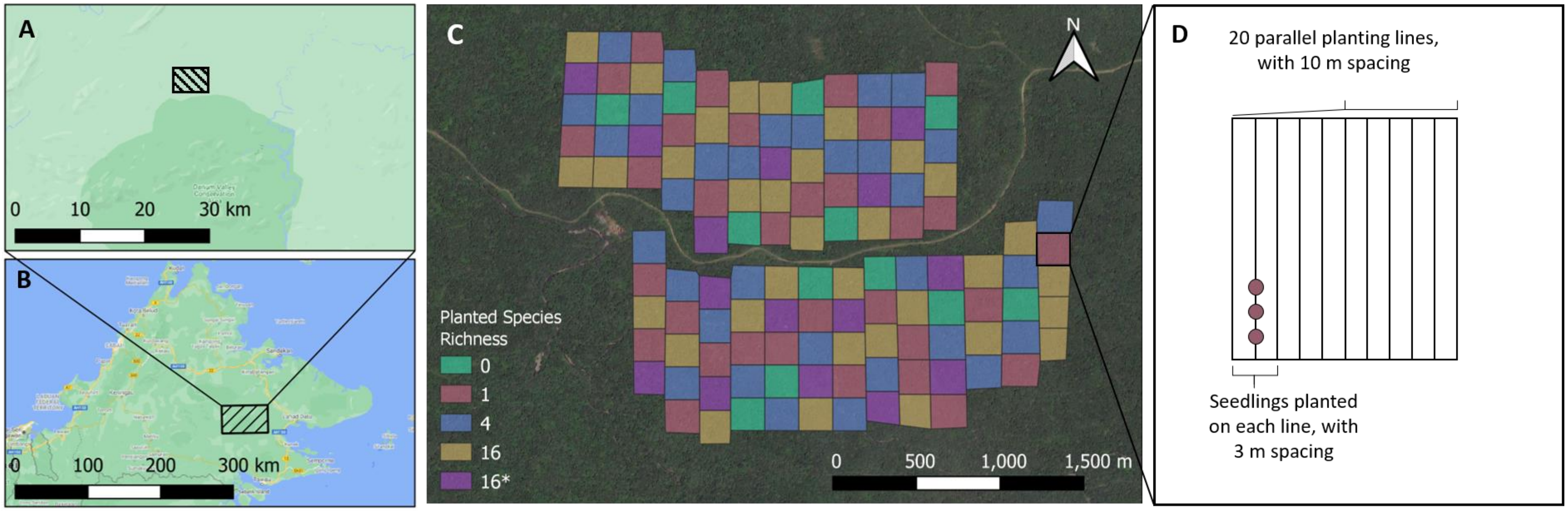


**Fig. S1. Location and design of the Sabah Biodiversity Experiment.** (**A** to **B**) The experimental site is located in a selectively logged area of the Malua Forest Reserve just north of the Danum Valley Conservation Area, Sabah, Malaysian Borneo. (**C**) The Sabah Biodiversity Experiment consists of 124 four-hectare plots, separated into two blocks by a logging road, with a combination of treatments that vary the species richness of enrichment-planted dipterocarp seedlings (0, 1, 4, or 16 species) and restoration methodologies (liana removal or not). (**D**) In each plot, dipterocarp seedlings were planted in transects 10 m apart, with 3 m spacing.

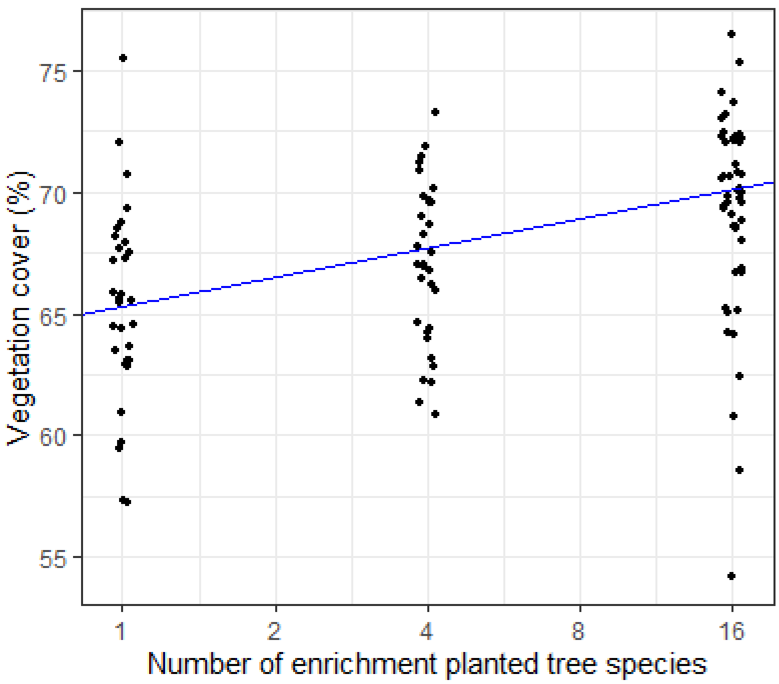


**Fig. S2. Effect of diversity on enrichment planted dipterocarp estimated vegetation cover.** Estimated vegetation cover (RapidEye) as a function of the number of enrichment-planted tree species a decade after initial planting. The line is the regression slope with the $\log_2$ number of tree species from the mixed-effects model analysis (points are jittered to avoid overlap).

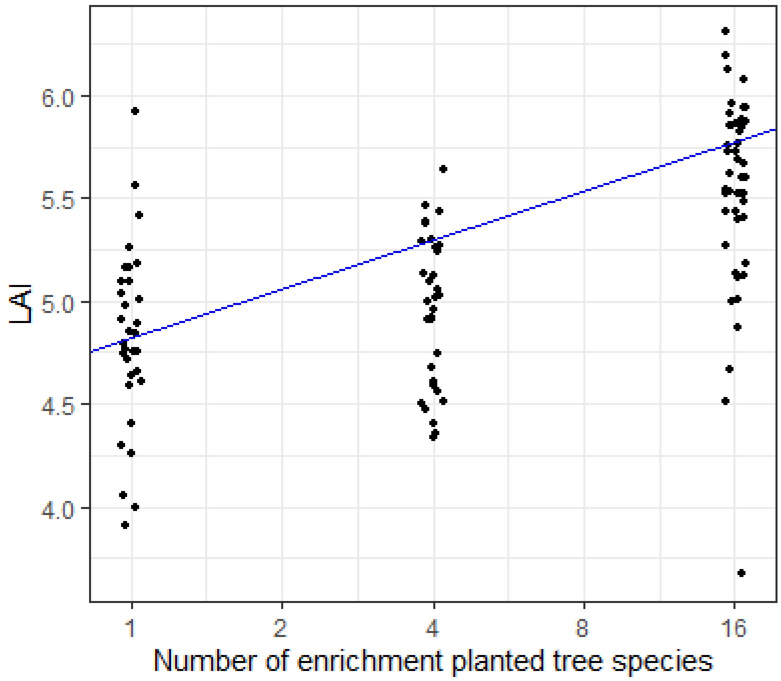


**Fig. S3. Effect of diversity on enrichment planted tree's estimated LAI.** Estimated LAI (RapidEye) as a function of the number of enrichment-planted tree species a decade after initial planting. The line is the regression slope with the $\log_2$ number of tree species from the mixed-effects model analysis (points are jittered to avoid overlap).

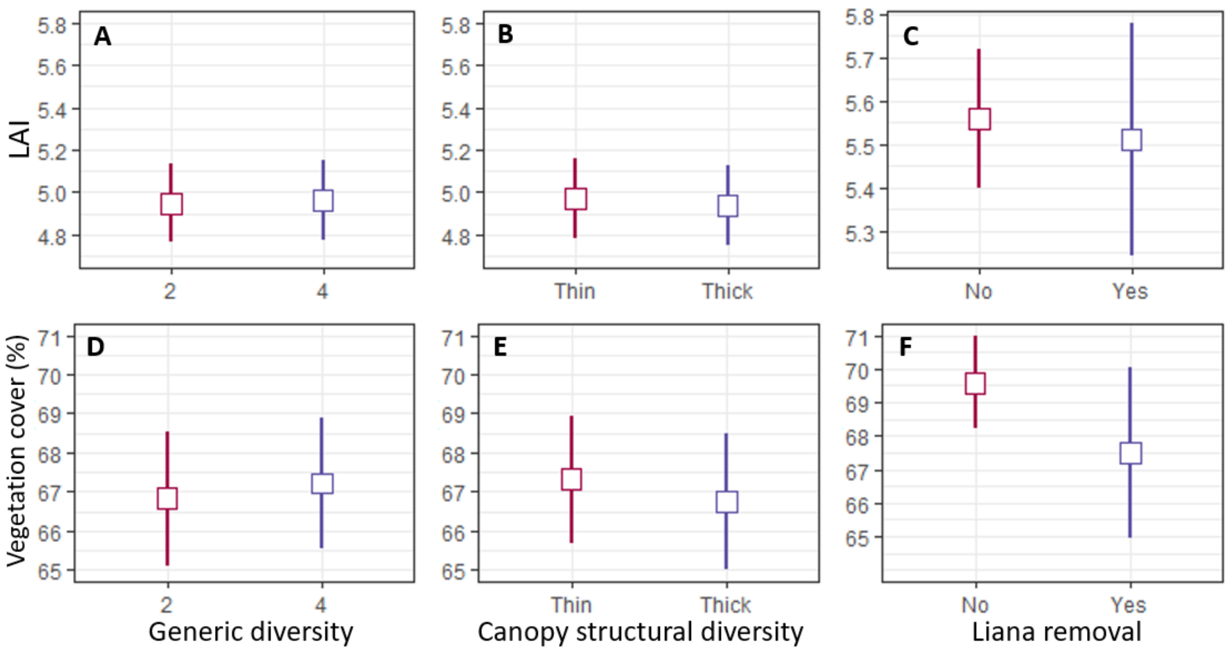


**Fig. S4. RapidEye satellite remote sensing estimates as a function of restoration treatment a decade after initial planting.** LAI (**A** to **C**) and cover (**D** to **F**) as a function of (from left to right) generic diversity of plots enrichment planted with four-species (2 genera vs 4 genera); canopy complexity with four species (low vs high); and liana ('climber') cutting with 16 species.

**Table S1. The 16 species of the Dipterocarpaceae family involved in the Sabah Biodiversity Experiment and relevant traits**

| Genus | Species | Species authority | Relative canopy height | Ecology | Timber group | IUCN status |
|---|---|---|---|---|---|---|
| Dipterocarpus | conformis | Slooten | Tall | Rare, Hill dipterocarp forest, clay rich soils, below 800 m | - | Endangered |
| Dryobalanops | lanceolata | Burck | Tall | Widespread on fertile soils, abundant on undulating land on volcanic/calcareous soils, up to 700 m | - | Least Concern |
| Hopea | ferruginea | Parijs | Short | Deep fertile soils in mixed dipterocarp forest, below 750 m | - | Critically endangered |
| | sangal | Korth. | Short | Often on or near riverbanks in low country and up to 500 m | - | Vulnerable |
| Parashorea | malaanonan | (Blanko) Merr. | Tall | Abundant in lowlands, typically E. Sabah, recorded up to 1300 m | - | Least concern |
| | tomentella | Meijer | Tall | Common on flat and undulating land, up to 200 m | - | Least concern |
| Shorea | argentifolia | Sym. | Medium | Locally frequent in forests, especially clay soils on undulating land and in valleys, below 600 m | Red Meranti | Least concern |
| | beccariana | Bruck | Medium | Common on leached lowland soils and dry ridges up to 1350 m | Red Meranti | Least Concern |
| | faguetiana | Heim. | Tall | Well-drained clay soils on low hills, and particularly ridge tops at 150-100 m (typically 700 m) | Yellow Meranti | Endangered |
| | gibbosa | Brandis. | Tall | Common on deep fertile soils, below 600 m | Yellow Meranti | Critically endangered |
| | johorensis | Foxw. | Tall | E. Borneo on well-drained fertile soils, below 600 m | Red Meranti | Critically endangered |
| | leprosula | Miq. | Medium | Deep clay soils in mixed dipterocarp forest below 700 m | Red Meranti | Near-threatened |
| | macrophylla | Ashton | Medium | Locally abundant on periodically flooded alluvium and riverbanks but rarer on hillsides, below 600 m | Red Meranti | Least Concern |
| | macroptera | King | Medium | Clay soils on low hills up to 600 m | Red Meranti | Least Concern |
| | ovalis | Korth. | Medium | Scattered in mixed dipterocarp forests, usually in moist or low-lying ground, up to 500 m | Red Meranti | Least Concern |
| | parvifolia | Dyer. | Medium | Perhaps the commonest dipterocarp, on clay soils on hills below 800 m | Red Meranti | Least Concern |

**Table S2. Mixed-effects model estimates of aboveground biomass, leaf area index and canopy cover for treatments**

| Treatment | Number of plots | AGB estimate (Mg $ha^{-1}$) | AGB SE | LAI Estimate | LAI SE | Cover estimate (%) | Cover SE |
|---|---|---|---|---|---|---|---|
| Unplanted | 12 | 182.67 | 4.26 | 4.57 | 0.246 | 62.05 | 2.28 |
| Monoculture | 32 | 214.89 | 3.15 | 4.82 | 0.231 | 65.33 | 2.15 |
| 4 species | 32 | 231.90 | 3.15 | 4.96 | 0.231 | 67.03 | 2.15 |
| 16 species | 38 | 261.51 | 3.15 | 5.53 | 0.231 | 69.34 | 2.15 |
| 16 species with liana removal | 10 | 264.18 | 3.86 | 5.64 | 0.240 | 69.31 | 2.23 |

**Table S3. Mixed-effects model estimates of aboveground biomass, leaf area index and canopy cover for unplanted vs planted plots**

| Treatment | Number of plots | AGB estimate (Mg ha$^{-1}$) | AGB SE | LAI estimate | LAI SE | Cover estimate (%) | Cover SE |
|---|---|---|---|---|---|---|---|
| Unplanted | 12 | 182.67 | 15.03 | 4.57 | 0.35 | 62.05 | 2.65 |
| Planted | 110 | 225.98. | 3.60 | 4.96 | 0.23 | 66.68 | 2.09 |

**Table S4. % cover recovery per doubling in species richness by epoch**

| Epoch | log2 canopy cover recovery rate estimate (% species richness$^{-1}$) | log2 canopy cover recovery rate SE |
|---|---|---|
| 1999-2002 | -0.0146 | 0.1415 |
| 2003-2008 | 0.256 | 0.0602 |
| 2008-2012 | 0.859 | 0.0602 |

**Table S5. 4-species mixture compositions**

**A) 2-genera diversity plots**

| | Thin structural diversity | | Thick structural diversity |
|---|---|---|---|
| **4.1** | *Parashorea malaanonan*<br>*Parashorea tomentella*<br>*Shorea beccariana*<br>*Shorea leprosula* | **4.5** | *Hopea sangal*<br>*Hopea ferruginea*<br>*Shorea beccariana*<br>*Shorea johorensis* |
| **4.2** | *Parashorea malaanonan*<br>*Parashorea tomentella*<br>*Shorea macroptera*<br>*Shorea ovalis* | **4.6** | *Hopea sangal*<br>*Hopea ferruginea*<br>*Shorea macroptera*<br>*Shorea gibbosa* |
| **4.3** | *Hopea sangal*<br>*Hopea ferruginea*<br>*Shorea macrophylla*<br>*Shorea parvifolia* | **4.7** | *Hopea sangal*<br>*Hopea ferruginea*<br>*Shorea macrophylla*<br>*Shorea faguetiana* |
| **4.4** | *Hopea sangal*<br>*Hopea ferruginea*<br>*Shorea argentifolia*<br>*Shorea parvifolia* | **4.8** | *Hopea sangal*<br>*Hopea ferruginea*<br>*Shorea argentifolia*<br>*Shorea johorensis* |

**B) 4-genera diversity plots**

| | Thin structural diversity | | Thick structural diversity |
|---|---|---|---|
| **4.9** | *Dipterocarpus conformis*<br>*Dryobalanops lanceolata*<br>*Parashorea malaanonan*<br>*Shorea faguetiana* | **4.13** | *Dipterocarpus conformis*<br>*Dryobalanops lanceolata*<br>*Shorea macrophylla*<br>*Hopea sangal* |
| **4.10** | *Dipterocarpus conformis*<br>*Dryobalanops lanceolata*<br>*Parashorea tomentella*<br>*Shorea johorensis* | **4.14** | *Dipterocarpus conformis*<br>*Dryobalanops lanceolata*<br>*Shorea ovalis*<br>*Hopea ferruginea* |
| **4.11** | *Dipterocarpus conformis*<br>*Dryobalanops lanceolata*<br>*Parashorea tomentella*<br>*Shorea gibbosa* | **4.15** | *Dipterocarpus conformis*<br>*Dryobalanops lanceolata*<br>*Shorea ovalis*<br>*Hopea sangal* |
| **4.12** | *Dipterocarpus conformis*<br>*Dryobalanops lanceolata*<br>*Parashorea malaanonan*<br>*Shorea johorensis* | **4.16** | *Dipterocarpus conformis*<br>*Dryobalanops lanceolata*<br>*Shorea macrophylla*<br>*Hopea ferruginea* |

**Table S6. AGB, LAI, and % cover estimates and SE by generic diversity and canopy structural diversity combinations, within 4-species plot combinations**

| | Canopy structural diversity | Number of plots | AGB estimate (Mg $ha^{-1}$) | AGB SE | LAI estimate | LAI SE | Cover estimate (%) | Cover SE |
|---|---|---|---|---|---|---|---|---|
| 2 genera | Thin | 8 | 231.0 | 3.67 | 5.02 | 0.246 | 67.4 | 2.19 |
| | Thick | 8 | 229.0 | 3.67 | 4.88 | 0.246 | 66.2 | 2.19 |
| 4 genera | Thin | 8 | 235.0 | 3.67 | 4.93 | 0.246 | 67.4 | 2.19 |
| | Thick | 8 | 232.0 | 3.67 | 5.00 | 0.246 | 67.2 | 2.19 |

**Table S7. AGB, LAI, and % cover estimates and SE for untreated and liana-removed 16-species plots**

| Liana removal | Number of plots | AGB estimate (Mg $ha^{-1}$) | AGB SE | LAI estimate | LAI SE | Cover estimate (%) | Cover SE |
|---|---|---|---|---|---|---|---|
| No | 38 | 261.51 | 3.15 | 5.53 | 0.231 | 69.34 | 2.15 |
| Yes | 10 | 264.18 | 3.86 | 5.64 | 0.240 | 69.31 | 2.23 |

**Table S8. Remote sensing datasets used in this study**

| Data | Spatial Resolution | Temporal Resolution | Time Cover |
|---|---|---|---|
| *Landsat VCF vegetation cover* | 30 m | 5-year | 2000, 2005, 2010* |
| *RapidEye-based vegetation cover* | 5 m | / | Aug. 2012 |
| *RapidEye-based leaf area index (LAI)* | 5 m | / | Aug. 2012 |
| *RapidEye-based aboveground biomass* | 5 m | / | Aug. 2012 |
| *MODIS MCD15A3H LAI* | 500 m | 4-day | 2001-2017 |

***Mid-points of the monitoring periods ('epochs’) – see text for details.**
