## Supplementary material for "Positive effects of tree diversity on tropical forest restoration in a field-scale experiment": Data analysis supplement

### Positive effects of tree diversity on tropical forest restoration in a field-scale experiment
### Supplementary material - analysis

Ryan Veryard

30 August, 2022

### 1 Introduction

This document accompanies the research article `Positive effects of tree diversity on tropical forest restoration in a field-scale experiment`, as part of a supplementary material. This document aims to provide an explanation of the analytical methodology used, and contains both the code as well as the outputs of each stage of the analysis, as well as an explanation as to why each stage was carried out. This document therefore aims to help ensure that this research is as reproducible as possible. This document will be present in both PDF and RMarkdown file types.

In our research article we investigated the effect of replanting, planted species richness, planted generic diversity and canopy complexity, and the effect of liana removal on growth within a secondary tropical forest in Sabah, Malaysian Borneo. For a detailed overview of the experimental design of this study, please see the main text together with the supplementary methods document.

#### 1.1 Approach to the analysis

Our analysis addresses the following questions:

1. Does replanting have an effect on forest biomass and structure (cover and LAI)?
2. Does the species richness of the enrichment planted seedlings impact forest biomass and structure?
3. Does the replanted generic diversity and canopy complexity impact forest biomass and structure?
4. Does the effect of additional climber cutting impact forest biomass and structure?

Our analysis was performed in R (version 3.6.3), using primarily the tidyverse (version 1.2.1) and lme4 (version 1.1-21) packages. Our analysis tries to address our a prior research questions (implicit in the experimental design) in as few models as possible utilizing a priori contrasts to address the comparisons of interest.

### 2 Read and Check Data

Read in the data files used in this analysis, and check that it is encoded correctly.

```
library(arm)
library(scales)
library(GGally)
library(lme4)
library(lmerTest)
library(RColorBrewer)
library(tidyverse)
library(plotrix)
library(FD)
library(cati)
library(picante)

select <- dplyr::select
theme_set(theme_bw())
```

```
sessionInfo()
```

```
## R version 4.1.0 (2021-05-18)
## Platform: x86_64-w64-mingw32/x64 (64-bit)
## Running under: Windows 10 x64 (build 17763)
##
## Matrix products: default
##
## locale:
## [1] LC_COLLATE=English_United Kingdom.1252
## [2] LC_CTYPE=English_United Kingdom.1252
## [3] LC_MONETARY=English_United Kingdom.1252
## [4] LC_NUMERIC=C
## [5] LC_TIME=English_United Kingdom.1252
##
## attached base packages:
## [1] stats     graphics  grDevices utils     datasets  methods   base
##
## other attached packages:
##  [1] picante_1.8.2      cati_0.99.3        nlme_3.1-152       FD_1.0-12
##  [5] vegan_2.5-7        lattice_0.20-44    permute_0.9-5      geometry_0.4.5
##  [9] ape_5.5            ade4_1.7-17        plotrix_3.8-1      forcats_0.5.1
## [13] stringr_1.4.0      dplyr_1.0.7        purrr_0.3.4        readr_2.0.1
## [17] tidyr_1.1.3        tibble_3.1.3       tidyverse_1.3.1    RColorBrewer_1.1-2
## [21] lmerTest_3.1-3     GGally_2.1.2       ggplot2_3.3.5      scales_1.1.1
## [25] arm_1.11-2         lme4_1.1-27.1      Matrix_1.3-3       MASS_7.3-54
##
## loaded via a namespace (and not attached):
##   [1] minqa_1.2.4         colorspace_2.0-2    ellipsis_0.3.2
##   [4] class_7.3-19        mclust_5.4.7        htmlTable_2.2.1
##   [7] base64enc_0.1-3     fs_1.5.0            rstudioapi_0.13
##  [10] proxy_0.4-26        hexbin_1.28.2       mvtnorm_1.1-2
##  [13] fansi_0.5.0         lubridate_1.8.0     xml2_1.3.2
##  [16] codetools_0.2-18    splines_4.1.0       knitr_1.33
```

```
##  [19] Formula_1.2-4       jsonlite_1.7.2      nloptr_1.2.2.2
##  [22] broom_0.7.9         cluster_2.1.2       dbplyr_2.1.1
##  [25] rgeos_0.5-5         png_0.1-7           compiler_4.1.0
##  [28] httr_1.4.2          backports_1.2.1     assertthat_0.2.1
##  [31] fastmap_1.1.0       cli_3.0.1           formatR_1.11
##  [34] prettyunits_1.1.1   htmltools_0.5.2     tools_4.1.0
##  [37] coda_0.19-4         gtable_0.3.0        glue_1.4.2
##  [40] maps_3.3.0          Rcpp_1.0.7          raster_3.4-13
##  [43] cellranger_1.1.0    vctrs_0.3.8         crosstalk_1.1.1
##  [46] fastcluster_1.2.3   xfun_0.25           rvest_1.0.1
##  [49] lifecycle_1.0.1     terra_1.3-4         zoo_1.8-9
##  [52] hms_1.1.0           parallel_4.1.0      yaml_2.2.1
##  [55] gridExtra_2.3       rpart_4.1-15        reshape_0.8.8
##  [58] latticeExtra_0.6-29 stringi_1.7.5       hypervolume_2.0.12
##  [61] e1071_1.7-8         checkmate_2.0.0     boot_1.3-28
##  [64] rlang_0.4.11        pkgconfig_2.0.3     rgl_0.107.10
##  [67] pracma_2.3.3        evaluate_0.14       ks_1.13.2
##  [70] htmlwidgets_1.5.3   pdist_1.2           tidyselect_1.1.1
##  [73] plyr_1.8.6          magrittr_2.0.1      R6_2.5.1
##  [76] generics_0.1.0      Hmisc_4.5-0         DBI_1.1.1
##  [79] pillar_1.6.2        haven_2.4.3         foreign_0.8-81
##  [82] withr_2.4.2         mgcv_1.8-35         sp_1.4-5
##  [85] survival_3.2-11     abind_1.4-5         nnet_7.3-16
##  [88] modelr_0.1.8        crayon_1.4.1        KernSmooth_2.23-20
##  [91] utf8_1.2.2          tzdb_0.1.2          rmarkdown_2.10
##  [94] progress_1.2.2      jpeg_0.1-9          grid_4.1.0
##  [97] readxl_1.3.1        data.table_1.14.0   reprex_2.0.1
## [100] digest_0.6.28       numDeriv_2016.8-1.1 rasterVis_0.50.3
## [103] munsell_0.5.0       viridisLite_0.4.0   magic_1.5-9
```

##### 2.1 Sat_data import

First the main dataset is loaded in, which contains the entirety of our data in a single large file.

```
Sat_data <- readRDS("Data/SBE_Data.rds")
```

```
str(Sat_data)
```

```
## tibble [744 x 17] (S3: tbl_df/tbl/data.frame)
##  $ Block          : Factor w/ 2 levels "North","South": 2 2 2 2 2 2 2 2 2 2 ...
##  $ Plot           : Factor w/ 124 levels "1","2","3","4",..: 1 1 1 1 1 1 6 6 6 6 ...
##  $ Spp_richness   : num [1:744] 4 4 4 4 4 4 4 4 4 4 ...
##  $ Spp_comp       : Factor w/ 35 levels "0_Unplanted",..: 24 24 24 24 24 24 32 32 32 32 ...
##  $ Climber_cutting: Factor w/ 2 levels "No","Yes": 1 1 1 1 1 1 1 1 1 1 ...
##  $ Satellite      : Factor w/ 2 levels "Landsat","RapidEye": 1 1 1 2 2 2 1 1 1 2 ...
##  $ Index          : Factor w/ 3 levels "Biomass","Cover",..: 2 2 2 3 1 2 2 2 2 3 ...
##  $ Year           : Factor w/ 3 levels "2000","2005",..: 1 2 3 NA NA NA 1 2 3 NA ...
##  $ Score          : num [1:744] 72.13 73.94 77.17 4.92 233.15 ...
##  $ Gen_div        : Factor w/ 4 levels "0","2","4","5": 3 3 3 3 3 3 2 2 2 2 ...
##  $ Canopy_type    : Factor w/ 3 levels "0","Thin","Thick": 3 3 3 3 3 3 3 3 3 3 ...
##  $ Canopy_str     : Factor w/ 7 levels "0","S","M","T",..: 7 7 7 7 7 7 7 7 7 7 ...
##  $ Planting       : Factor w/ 2 levels "Control","Planted": 2 2 2 2 2 2 2 2 2 2 ...
```

```
##  $ Treatment      : Factor w/ 5 levels "0_spp","1_spp",..: 3 3 3 3 3 3 3 3 3 3 ...
##  $ Random         : Factor w/ 2 levels "No","Yes": 2 2 2 2 2 2 2 2 2 2 ...
##  $ PD             : num [1:744] 0.0655 0.0655 0.0655 0.0655 0.0655 ...
##  $ FD             : num [1:744] 2.2 2.2 2.2 2.2 2.2 ...
```

```
summary(Sat_data)
```

```
##    Block          Plot      Spp_richness                          Spp_comp
##  North:360   1       :  6   Min.   : 0.000   16_spp                   :228
##  South:384   2       :  6   1st Qu.: 1.000   0_Unplanted              : 72
##              3       :  6   Median : 4.000   16_cut                   : 60
##              4       :  6   Mean   : 7.484   1_Dipterocarpus_conformis: 12
##              5       :  6   3rd Qu.:16.000   1_Dryobalanops_lanceolate: 12
##              6       :  6   Max.   :16.000   1_Hopea_ferruginea       : 12
##              (Other):708                     (Other)                  :348
##  Climber_cutting    Satellite       Index        Year         Score
##  No :684         Landsat :372   Biomass:124   2000:124   Min.   :  3.671
##  Yes: 60         RapidEye:372   Cover  :496   2005:124   1st Qu.: 67.436
##                                 LAI    :124   2010:124   Median : 73.481
##                                               NA's:372   Mean   : 88.475
##                                                          3rd Qu.: 77.374
##                                                          Max.   :290.831
##
##  Gen_div Canopy_type Canopy_str    Planting     Treatment    Random
##  0: 72   0    : 72   0  : 72    Control: 72   0_spp : 72   No :132
##  2:288   Thin :288   S  : 24    Planted:672   1_spp :192   Yes:612
##  4: 96   Thick:384   M  : 84                  4_spp :192
##  5:288               T  :120                  16_spp:228
##                      SM : 24                  16_cut: 60
##                      MT : 36
##                      SMT:384
##        PD                 FD
##  Min.   :0.01467   Min.   :0.000
##  1st Qu.:0.01846   1st Qu.:0.000
##  Median :0.05935   Median :2.095
##  Mean   :0.05880   Mean   :1.932
##  3rd Qu.:0.09438   3rd Qu.:3.594
##  Max.   :0.09438   Max.   :3.594
##  NA's   :72
```

##### 2.2 Rapideye_data and Landsat_data import

For ease of some of the analyses contained within this document, the original `Sat_data` dataset has been divided into two smaller datasets, `Rapideye_data` and `Landsat_data`, representing data from the RapidEye and Landsat Satellites. More detail of how these two dataset structures is explained below.

```
Rapideye_data <- readRDS("Data/RapidEye_SBE_Data.rds")
```

```
str(Rapideye_data)
```

```
## tibble [124 x 16] (S3: tbl_df/tbl/data.frame)
```

```
##  $ Block          : Factor w/ 2 levels "North","South": 2 2 2 2 2 2 2 2 2 2 ...
##  $ Plot           : Factor w/ 124 levels "1","2","3","4",..: 1 6 9 12 16 19 25 28 35 39 ...
##  $ Spp_richness   : num [1:124] 4 4 4 4 4 4 4 4 4 4 ...
##  $ Spp_comp       : Factor w/ 35 levels "0_Unplanted",..: 24 32 29 21 22 31 28 33 23 35 ...
##  $ Climber_cutting: Factor w/ 2 levels "No","Yes": 1 1 1 1 1 1 1 1 1 1 ...
##  $ Gen_div        : Factor w/ 4 levels "0","2","4","5": 3 2 2 3 3 2 2 2 3 3 ...
##  $ Canopy_type    : Factor w/ 3 levels "0","Thin","Thick": 3 3 2 2 2 3 2 3 2 2 ...
##  $ Canopy_str     : Factor w/ 7 levels "0","S","M","T",..: 7 7 5 4 4 7 6 7 6 4 ...
##  $ Planting       : Factor w/ 2 levels "Control","Planted": 2 2 2 2 2 2 2 2 2 2 ...
##  $ Treatment      : Factor w/ 5 levels "0_spp","1_spp",..: 3 3 3 3 3 3 3 3 3 3 ...
##  $ Random         : Factor w/ 2 levels "No","Yes": 2 2 2 2 2 2 2 2 2 2 ...
##  $ PD             : num [1:124] 0.0655 0.0404 0.0371 0.0564 0.0574 ...
##  $ FD             : num [1:124] 2.2 2.05 2.03 2.14 2.17 ...
##  $ LAI            : num [1:124] 4.92 5.27 4.96 4.59 5.09 ...
##  $ Biomass        : num [1:124] 233 234 226 247 242 ...
##  $ Cover          : num [1:124] 66.4 69.6 67 66.2 68.3 ...
```

```
summary(Rapideye_data)
```

```
##    Block         Plot      Spp_richness                         Spp_comp
##  North:60   1      :  1   Min.   : 0.000   16_spp                   :38
##  South:64   2      :  1   1st Qu.: 1.000   0_Unplanted              :12
##             3      :  1   Median : 4.000   16_cut                   :10
##             4      :  1   Mean   : 7.484   1_Dipterocarpus_conformis: 2
##             5      :  1   3rd Qu.:16.000   1_Dryobalanops_lanceolate: 2
##             6      :  1   Max.   :16.000   1_Hopea_ferruginea       : 2
##             (Other):118                    (Other)                  :58
##  Climber_cutting Gen_div Canopy_type Canopy_str    Planting    Treatment
##  No :114         0:12    0    :12    0  :12     Control: 12   0_spp :12
##  Yes: 10         2:48    Thin :48    S  : 4     Planted:112   1_spp :32
##                  4:16    Thick:64    M  :14                   4_spp :32
##                  5:48                T  :20                   16_spp:38
##                                      SM : 4                   16_cut:10
##                                      MT : 6
##                                      SMT:64
##  Random          PD                FD              LAI           Biomass
##  No : 22   Min.   :0.01467   Min.   :0.000   Min.   :3.671   Min.   :167.4
##  Yes:102   1st Qu.:0.01846   1st Qu.:0.000   1st Qu.:4.704   1st Qu.:215.2
##            Median :0.05935   Median :2.095   Median :5.103   Median :234.7
##            Mean   :0.05880   Mean   :1.932   Mean   :5.114   Mean   :234.2
##            3rd Qu.:0.09438   3rd Qu.:3.594   3rd Qu.:5.528   3rd Qu.:259.6
##            Max.   :0.09438   Max.   :3.594   Max.   :6.308   Max.   :290.8
##            NA's   :12
##      Cover
##  Min.   :54.16
##  1st Qu.:64.19
##  Median :67.37
##  Mean   :66.93
##  3rd Qu.:70.06
##  Max.   :76.45
##
```

```r
Landsat_data <- readRDS("Data/Landsat_SBE_Data.rds")
```

```r
str(Landsat_data)
```

```
## tibble [372 x 15] (S3: tbl_df/tbl/data.frame)
##  $ Block          : Factor w/ 2 levels "North","South": 2 2 2 2 2 2 2 2 2 2 ...
##  $ Plot           : Factor w/ 124 levels "1","2","3","4",..: 1 1 1 6 6 6 9 9 9 12 ...
##  $ Spp_richness   : num [1:372] 4 4 4 4 4 4 4 4 4 4 ...
##  $ Spp_comp       : Factor w/ 35 levels "0_Unplanted",..: 24 24 24 32 32 32 29 29 29 21 ...
##  $ Climber_cutting: Factor w/ 2 levels "No","Yes": 1 1 1 1 1 1 1 1 1 1 ...
##  $ Year           : Factor w/ 3 levels "2000","2005",..: 1 2 3 1 2 3 1 2 3 1 ...
##  $ Cover          : num [1:372] 72.1 73.9 77.2 73.5 77.4 ...
##  $ Gen_div        : Factor w/ 4 levels "0","2","4","5": 3 3 3 2 2 2 2 2 2 3 ...
##  $ Canopy_type    : Factor w/ 3 levels "0","Thin","Thick": 3 3 3 3 3 3 2 2 2 2 ...
##  $ Canopy_str     : Factor w/ 7 levels "0","S","M","T",..: 7 7 7 7 7 7 5 5 5 4 ...
##  $ Planting       : Factor w/ 2 levels "Control","Planted": 2 2 2 2 2 2 2 2 2 2 ...
##  $ Treatment      : Factor w/ 5 levels "0_spp","1_spp",..: 3 3 3 3 3 3 3 3 3 3 ...
##  $ Random         : Factor w/ 2 levels "No","Yes": 2 2 2 2 2 2 2 2 2 2 ...
##  $ PD             : num [1:372] 0.0655 0.0655 0.0655 0.0404 0.0404 ...
##  $ FD             : num [1:372] 2.2 2.2 2.2 2.05 2.05 ...
```

```r
summary(Landsat_data)
```

```
##    Block          Plot      Spp_richness                          Spp_comp
##  North:180   1      :  3   Min.   : 0.000   16_spp                    :114
##  South:192   2      :  3   1st Qu.: 1.000   0_Unplanted               : 36
##              3      :  3   Median : 4.000   16_cut                    : 30
##              4      :  3   Mean   : 7.484   1_Dipterocarpus_conformis:  6
##              5      :  3   3rd Qu.:16.000   1_Dryobalanops_lanceolate:  6
##              6      :  3   Max.   :16.000   1_Hopea_ferruginea       :  6
##              (Other):354                    (Other)                   :174
##  Climber_cutting   Year          Cover        Gen_div Canopy_type Canopy_str
##  No :342         2000:124   Min.   :68.77   0: 36   0    : 36   0  : 36
##  Yes: 30         2005:124   1st Qu.:72.87   2:144   Thin :144   S  : 12
##                  2010:124   Median :75.06   4: 48   Thick:192   M  : 42
##                             Mean   :74.86   5:144               T  : 60
##                             3rd Qu.:76.71                       SM : 12
##                             Max.   :81.45                       MT : 18
##                                                                 SMT:192
##     Planting      Treatment    Random         PD                FD
##  Control: 36   0_spp : 36   No : 66   Min.   :0.01467   Min.   :0.000
##  Planted:336   1_spp : 96   Yes:306   1st Qu.:0.01846   1st Qu.:0.000
##                4_spp : 96             Median :0.05935   Median :2.095
##                16_spp:114             Mean   :0.05880   Mean   :1.932
##                16_cut: 30             3rd Qu.:0.09438   3rd Qu.:3.594
##                                       Max.   :0.09438   Max.   :3.594
##                                       NA's   :36
```

##### 2.3 Descriptive summary of data

The structure of the `Sat_data` dataset incorporates the key elements of the Sabah Biodiversity Experiment Study Design:

- Two blocks (north and south), separated by an old logging road [Block]
- 124 plots, each 4-ha (200 x 200 m) in size [Plot]
- Each plot's associated planted species richness (between 0 (unplanted) and a 16-species mixture) [Spp_richness]
- Each plot's generic diversity, ranging between 0 and 5 [Gen_div]
  - The focus here is on 4-species mixtures, which were either planted with a generic diversity of 2 or 4. There is no variation within any other level of species richness
- The level of canopy complexity of the plot planted, measured in two ways:
  - Simple thick or thin combination of tree canopies [Canopy_type]
  - A more comprehensive breakdown of if the plot contains species with typically short, medium, and/or tall canopies [Canopy_str]
- If enhanced climber cutting was performed (a common management strategy in Sabah) [Climber_cutting]
  - Note: this treatment was only applied to a subset of the 16-species mixtures. Due to practical constraints, only the southern block had climber cutting treatments applied randomly, with northern block treatments determined by proximity to the road. As such an additional factor, Random, specifies if the treatment applies to each plot was done so randomly or not
- All 5 possible treatments (unplanted controls, monocultures, 4-species polycultures, 16-species polycultures, and 16-species polycultures with enhanced climber cutting) are captured in the factor Treatment

For each plot six measurements are taken;

- Three Landsat vegetation cover measurements, covering each of the three time periods covered ('epochs')
- One RapidEye measurement each for vegetation cover, LAI, and estimated biomass

For ease of analysis, Landsat and RapidEye datasets were separated into different datasets. Landsat vegetation cover scores were recorded in the column Score within the Landsat_data dataset, and Year specifies each epoch, encoded as a factor. Within the Rapideye_data dataset a separate response column stores each of the Biomass, Cover, and LAI values.

Finally, each dataset contains columns representing calculated phylogenetic and functional diversity measures for each plot's planted species richness:

- Faith's PD [PD]
- Functional diversity (as calculated by Petchy & Gaston (2002)) [FD]

A descriptive summary of the major differences between RapidEye and Landsat data can be seen below:

| Data | Spatial Resolution | Temporal Resolution | Time Cover |
|---|---|---|---|
| *Landsat VCF vegetation cover* | 30 m | 5-year | 2000, 2005, 2010 |
| *RapidEye-based vegetation cover* | 5 m | / | Aug. 2012 |
| *RapidEye-based leaf area index (LAI)* | 5 m | / | Aug. 2012 |
| RapidEye-based aboveground biomass | 5 m | / | Aug. 2012 |

We have the correct number of treatment groups, as specified in experimental design. Although we expect 16 plots of the treatment type `16_cut`, since the 6 plots within the southern block were *de facto* unplanted during August 2012 (when the RapidEye imagery was taken), those plots are classified here as `16_spp`.

```
Rapideye_data %>%
    group_by(Treatment) %>%
    summarise(n())
```

```
## # A tibble: 5 x 2
##   Treatment `n()`
##   <fct>     <int>
## 1 0_spp        12
## 2 1_spp        32
## 3 4_spp        32
## 4 16_spp       38
## 5 16_cut       10
```

We obtain, as expected, three times this in the `Landsat_data` dataset and six times these values in the `Sat_data` dataset.

```
Landsat_data %>%
    group_by(Treatment) %>%
    summarise(n())
```

```
## # A tibble: 5 x 2
##   Treatment `n()`
##   <fct>     <int>
## 1 0_spp        36
## 2 1_spp        96
## 3 4_spp        96
## 4 16_spp      114
## 5 16_cut       30
```

```
Sat_data %>%
    group_by(Treatment) %>%
    summarise(n())
```

```
## # A tibble: 5 x 2
##   Treatment `n()`
##   <fct>     <int>
## 1 0_spp        72
## 2 1_spp       192
## 3 4_spp       192
## 4 16_spp      228
## 5 16_cut       60
```

There are no inaccuracies with labelling between species composition, expected species richness, and planted vs controlled plots. This is correct for all three datasets:

```
Rapideye_data %>%
    group_by(Spp_comp, Spp_richness, Planting) %>%
    summarise(n())
```

```
## # A tibble: 35 x 4
## # Groups:   Spp_comp, Spp_richness [35]
##    Spp_comp                  Spp_richness Planting `n()`
##    <fct>                            <dbl> <fct>    <int>
##  1 0_Unplanted                          0 Control     12
##  2 1_Dipterocarpus_conformis            1 Planted      2
##  3 1_Dryobalanops_lanceolate            1 Planted      2
##  4 1_Hopea_ferruginea                   1 Planted      2
##  5 1_Hopea_sangal                       1 Planted      2
##  6 1_Parashorea_malaanonan              1 Planted      2
##  7 1_Parashorea_tomentalla              1 Planted      2
##  8 1_Shorea_argentifolia                1 Planted      2
##  9 1_Shorea_beccariana                  1 Planted      2
## 10 1_Shorea_faguetiana                  1 Planted      2
## # ... with 25 more rows
```

```
Landsat_data %>%
    group_by(Spp_comp, Spp_richness, Planting) %>%
    summarise(n())
```

```
## # A tibble: 35 x 4
## # Groups:   Spp_comp, Spp_richness [35]
##    Spp_comp                  Spp_richness Planting `n()`
##    <fct>                            <dbl> <fct>    <int>
##  1 0_Unplanted                          0 Control     36
##  2 1_Dipterocarpus_conformis            1 Planted      6
##  3 1_Dryobalanops_lanceolate            1 Planted      6
##  4 1_Hopea_ferruginea                   1 Planted      6
##  5 1_Hopea_sangal                       1 Planted      6
##  6 1_Parashorea_malaanonan              1 Planted      6
##  7 1_Parashorea_tomentalla              1 Planted      6
##  8 1_Shorea_argentifolia                1 Planted      6
##  9 1_Shorea_beccariana                  1 Planted      6
## 10 1_Shorea_faguetiana                  1 Planted      6
## # ... with 25 more rows
```

```
Sat_data %>%
    group_by(Spp_comp, Spp_richness, Planting) %>%
    summarise(n())
```

```
## # A tibble: 35 x 4
## # Groups:   Spp_comp, Spp_richness [35]
##    Spp_comp                  Spp_richness Planting `n()`
##    <fct>                            <dbl> <fct>    <int>
##  1 0_Unplanted                          0 Control     72
##  2 1_Dipterocarpus_conformis            1 Planted     12
##  3 1_Dryobalanops_lanceolate            1 Planted     12
##  4 1_Hopea_ferruginea                   1 Planted     12
##  5 1_Hopea_sangal                       1 Planted     12
##  6 1_Parashorea_malaanonan              1 Planted     12
##  7 1_Parashorea_tomentalla              1 Planted     12
##  8 1_Shorea_argentifolia                1 Planted     12
##  9 1_Shorea_beccariana                  1 Planted     12
```

```
## 10 1_Shorea_faguetiana                    1 Planted     12
## # ... with 25 more rows
```

### 3 Exploratory plotting

#### 3.1 Overall treatments

Observing the distribution of our RapidEye data (Fig.1) for each of our five treatments (each level of species richness as well as the 16-species mixture with climber cutting) we see across each of our indices that data is relatively symmetrical for each treatment, with only a small handful of outlying data points. `Cover` and `LAI` appear to be more similar to each other in distribution than to `Biomass`, which is expected due to their index calculation being more similar. `Biomass` scores are much more condensed, with a tighter distribution around their median values, and less overlap of violins than either `Cover` or `LAI`.

```
temp <- RColorBrewer::brewer.pal(5, "Spectral")
temp2 <- c("#a56327", "#dfc27d", "#8dc476", "#80cdc1",
    "#0d8672")

treatment_labels <- c("0", "1", "4", "16", "16*")
bio_label <- expression(paste("Aboveground Biomass (Mg ",
    Ha^-1, ")"))

Sat_data %>%
    filter(Satellite == "RapidEye") %>%
    ggplot(aes(Treatment, Score)) + geom_violin(aes(fill = Treatment),
    alpha = 0.6) + facet_wrap(~Index, scales = "free",
    strip.position = "left", labeller = as_labeller(c(Biomass = "Aboveground Biomass (Mg ha^-1)",
        LAI = "Leaf Area Index", Cover = "Cover (%)"))) +
    ylab(NULL) + theme(strip.background = element_blank(),
    strip.placement = "outside", strip.text = element_text(size = 11)) +
    geom_jitter(shape = 16, size = 0.9, position = position_jitter(0.1)) +
    scale_fill_manual(values = temp) + scale_x_discrete(labels = treatment_labels) +
    labs(x = "Number of Enrichment Planted Tree Species") +
    theme(legend.position = "none")
```

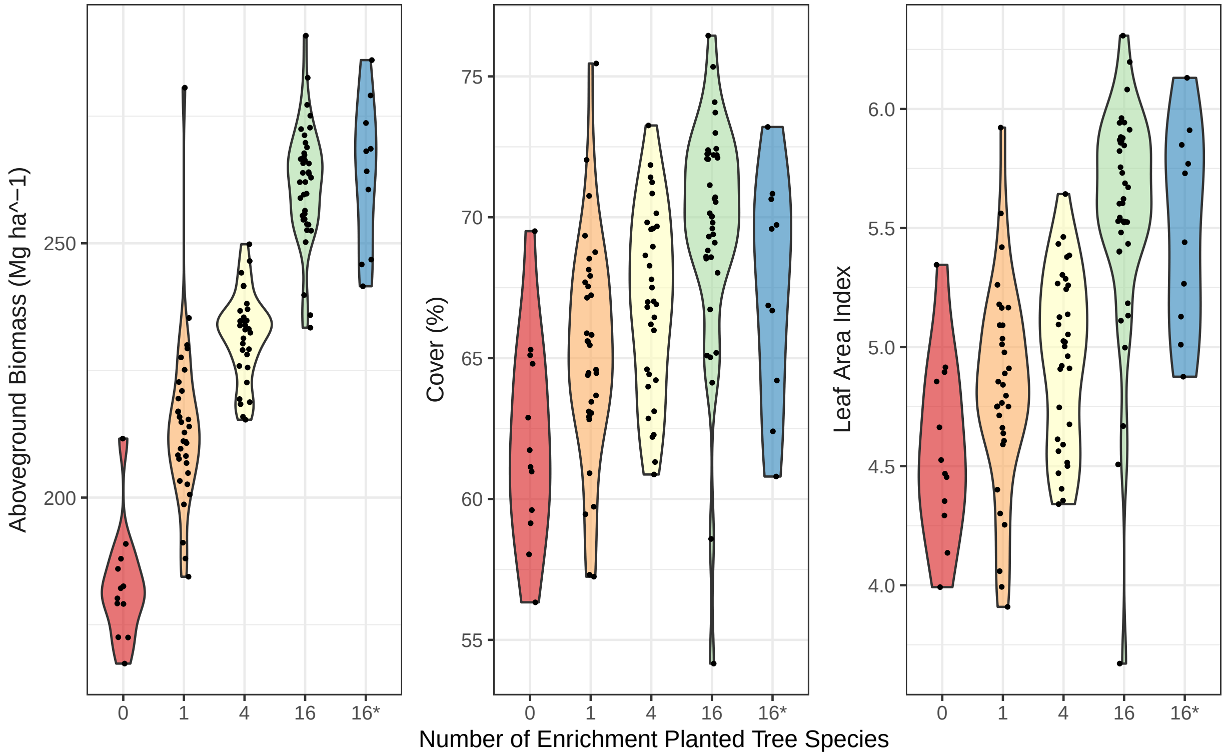

Figure 1: Data points for experimental plots overlaid on violin plots showing (left to right) aboveground biomass, percent vegetation cover, and Leaf Area Index in relation to enrichment planting with seedlings of 0, 1, 4, or 16 species of dipterocarp tree species (16*: enrichment planting with sixteen species plus liana cutting). Note: this figure relates to Fig. 1 (A to C) within the main manuscript.

#### 3.2 Covariation between measurements

We would expect measurement values for vegetation cover from both Landsat and RapidEye satellites to covary, as they are effectively trying to estimate the same values using different technology. Further, we have Landsat cover scores for three epochs (2000, 2005, and 2010). Although they are taken years apart, each pixel is more likely to be similar to itself across multiple years than to other index measurements (as it is the same calculation being carried out), so we should expect these values to covary too. Finally, of the three RapidEye indices (LAI, cover, and aboveground biomass), these should be expected to covary, since if cover was increasing we would expect the total amount of biomass to increase too, as well as the total leaf area coverage. This is what we see to be the case. The only index which does significantly correlate with other index values is the Landsat data from the 2000 epoch. These were measured at the very beginning of the experiment, whilst other values were measured 5-12 years later and after varied treatments being applied. This is therefore an understandable result.

Analysing every response variable at each stage of our analysis has drawbacks from a statistical perspective (including performing a large number of tests). To reduce the number of tests performed, it was chosen to continue analysis with RapidEye aboveground biomass response data. This is as it was based on images of higher resolution, and also was most relevant to the current literature.

```
pairs_data <- Sat_data %>%
    pivot_wider(names_from = c(Satellite, Index, Year),
        values_from = Score) %>%
    select(-c(Block:FD))
```

```r
colnames(pairs_data) <- gsub("_NA", "", colnames(pairs_data))

ggpairs(pairs_data)
```

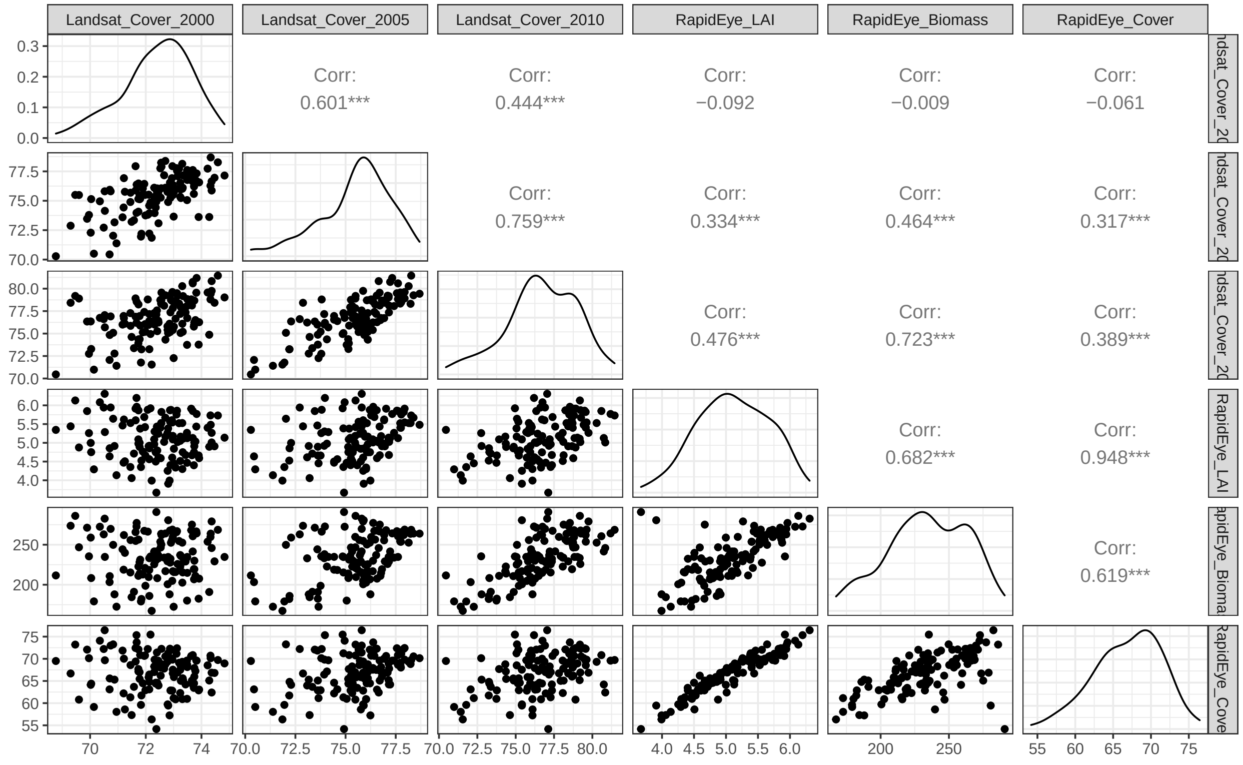

Figure 2: correlation plots for each satellite index.

##### 3.3 Effect of planting

A plot of the mean and standard error for plot AGB, cover, and LAI shows visually that there is some strong difference between the 12 unplanted controls at the SBE compared to all other planted sites, averaged across all other treatment types.

```r
cols <- RColorBrewer::brewer.pal(5, "Set1")
cols_2 <- c(brewer.pal(11, "Spectral")[1], brewer.pal(11,
    "Spectral")[11])
```

```r
temp <- Rapideye_data %>%
    pivot_longer(cols = c(Biomass, Cover, LAI), names_to = "Index",
        values_to = "Value") %>%
    group_by(Planting, Index) %>%
    summarise(Mean_Value = mean(Value), Se_Value = std.error(Value))

temp$Index = factor(temp$Index, labels = c("atop(Aboveground~Biomass,(Mg~ha^{-1}))",
    "Cover~('%')", "Leaf~Area~Index"))
```

```
ggplot(temp) + geom_pointrange(aes(x = Planting, y = Mean_Value,
    ymin = Mean_Value - 2 * Se_Value, ymax = Mean_Value +
        2 * Se_Value, colour = Planting), size = 1, fill = "white",
    shape = 22) + facet_wrap(~Index, scales = "free_y",
    strip.position = "left", labeller = label_parsed) +
    scale_colour_manual(values = c(cols_2[1], cols_2[2])) +
    theme(legend.position = "none") + labs(x = "Replanting treatment",
    y = NULL) + theme(strip.background = element_blank(),
    strip.placement = "outside", strip.text = element_text(size = 11),
    panel.spacing = unit(0, "lines"))
```

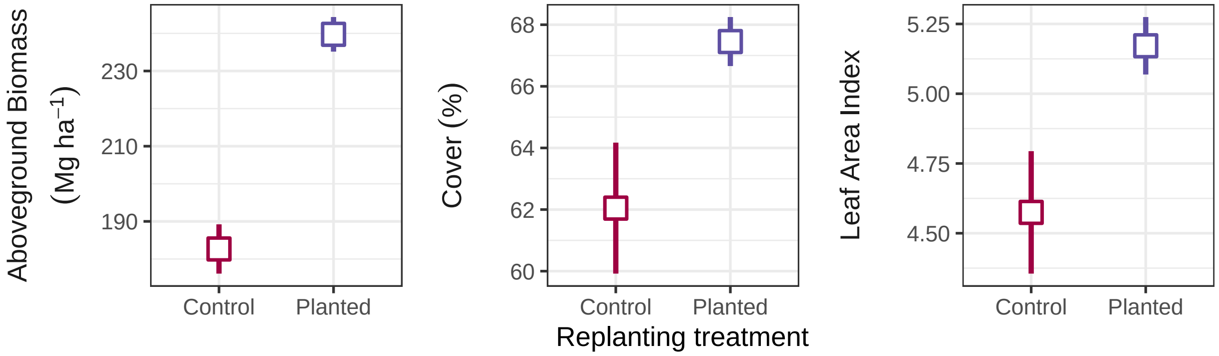

Figure 3: Treatment means (with 95% confidence intervals) for unplanted controls versus enrichment planted plots. Note: this figure relates to Fig. 1 (D to F) within the main manuscript.

#### 3.4 Effect of species richness

From the Landsat data, there seems to be a clear effect species richness on AGB estimates. As in many other biodiversity experiments, the relationship is approximately linear when species richness is log transformed (using base-2 so that each unit of the X axis corresponds to the response for every doubling of species richness):

```
Rapideye_data %>%
    filter(Spp_richness > 0) %>%
    ggplot(aes(Spp_richness, Biomass)) + geom_jitter(width = 0.07,
    size = 0.8) + geom_abline(slope = 13.3, colour = "blue",
    intercept = 213.41) + labs(x = "Number of enrichment planted tree species",
    y = expression(paste("Aboveground biomass (Mg  ", Ha^-1,
        ")"))) + scale_x_continuous(trans = log2_trans())
```

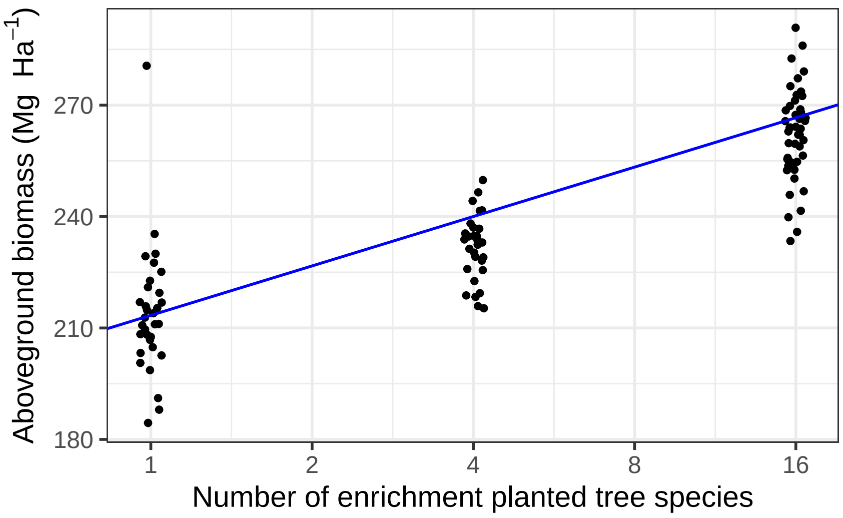

Figure 4: Estimated aboveground biomass as a function of the number of enrichment-planted tree species a decade after initial planting. Note: this figure relates to Fig. 2 (A) within the main manuscript.

Introducing the effect of time into these graphs, through the Landsat dataset, appears to show that different effects become clear at different stages of the experiment. Before the experiment takes place there is no difference between any level of species richness, but after ~ 5 years an effect of planting can be seen. Between 5-10 years an effect of species richness develops:

```
dodge <- position_dodge(0.3)
year_labels <- c("1999-2002", "2003-2008", "2008-2012")

Landsat_data %>%
    group_by(Year, Spp_richness) %>%
    summarise(mean_cover = mean(Cover), se_cover = std.error(Cover)) %>%
    ggplot(aes(Year, mean_cover, colour = as.factor(Spp_richness),
        group = as.factor(Spp_richness))) + geom_line(position = dodge,
    aes(linetype = as.factor(Spp_richness))) + geom_pointrange(aes(x = Year,
    y = mean_cover, ymin = mean_cover - 2 * se_cover, ymax = mean_cover +
        2 * se_cover, shape = as.factor(Spp_richness)),
    position = dodge, fill = "white", ) + scale_shape_manual(values = c(21:24)) +
    scale_colour_manual(values = cols) + scale_x_discrete(labels = year_labels) +
    labs(x = "Epoch", y = "Vegetation cover (%)", col = "Tree planted \nspecies richness",
        shape = "Tree planted \nspecies richness", linetype = "Tree planted \nspecies richness")
```

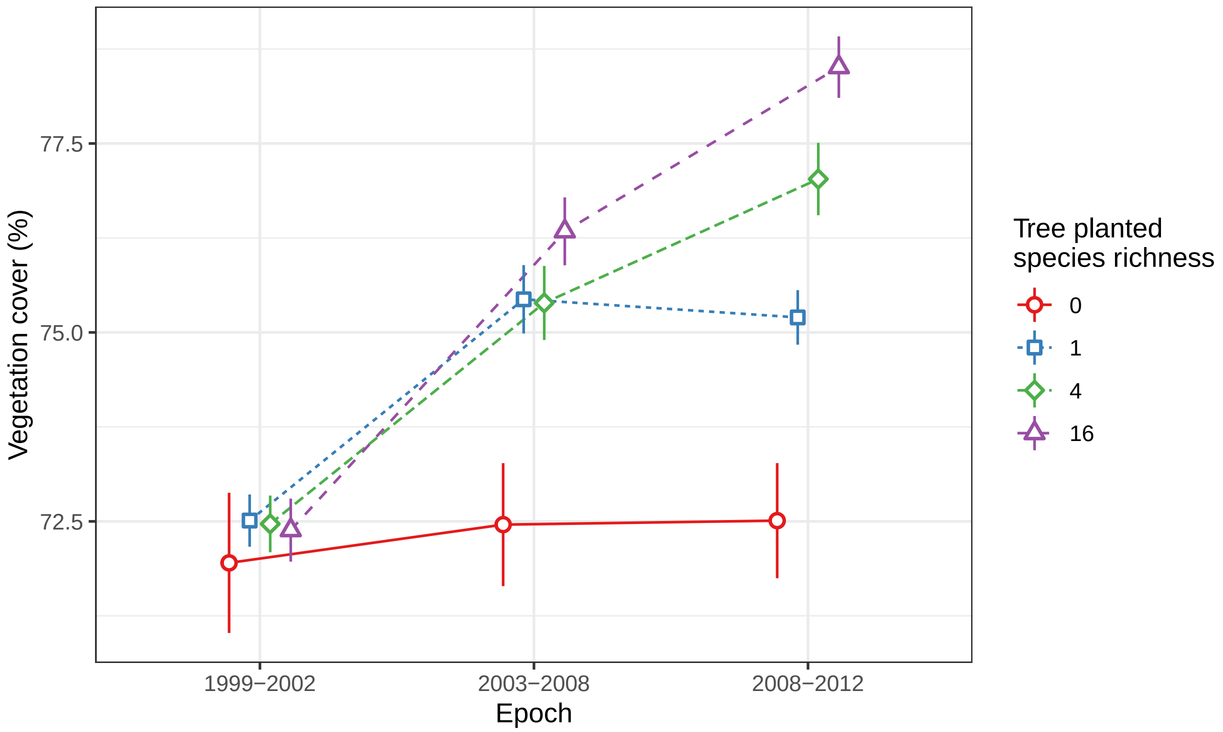

Figure 5: Changes in vegetation cover over time as a function of the number of enrichment-planted tree species. Note: this figure relates to Fig. 2 (B) within the main manuscript.

##### 3.5 Effect of planted generic diversity within 4-species mixtures

Although there is a greater mean AGB estimate with a generic diversity of 4 rather than 2, the effect seems to be far less than the variance associated with each group. It seems clear here, when holding species richness constant, there is no effect of planted generic diversity to be seen in the 2012 RapidEye imagery.

```
Rapideye_data %>%
    filter(Spp_richness == 4) %>%
    group_by(Gen_div) %>%
    summarise(Mean_Biomass = mean(Biomass), Se_Biomass = std.error(Biomass)) %>%
    ggplot(aes(x = Gen_div, y = Mean_Biomass)) + geom_pointrange(aes(ymin = Mean_Biomass -
    2 * Se_Biomass, ymax = Mean_Biomass + 2 * Se_Biomass,
    colour = Gen_div, shape = Gen_div), size = 1, fill = "white",
    shape = 22) + scale_colour_manual(values = cols_2) +
    theme(legend.position = "none") + labs(y = expression(atop("Aboveground biomass",
    paste("(Mg  ", Ha^-1, ")"))), x = "Generic diversity") +
    scale_y_continuous(limits = c(225, 239), breaks = c(225,
        230, 235))
```

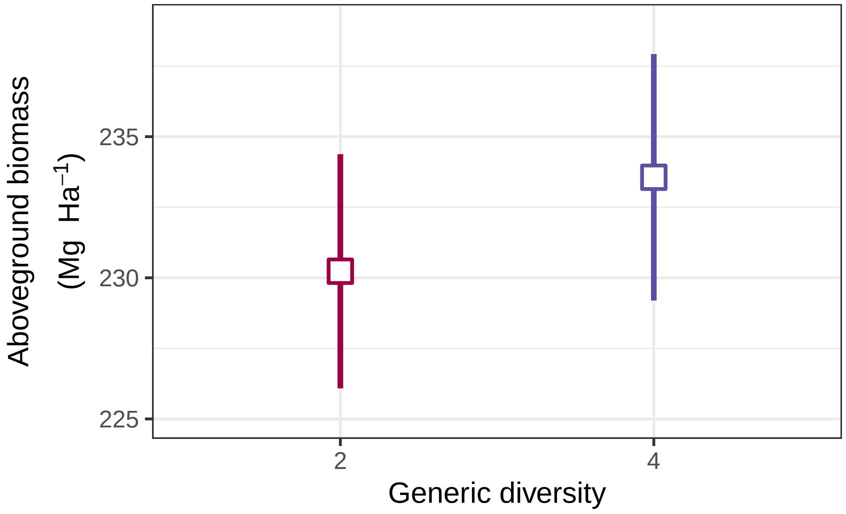

Figure 6: Aboveground biomass as a function of generic diversity of plots with four-species (2 vs 4 species). Note: this figure relates to Fig. 1 (I) within the main manuscript.

##### 3.6 Effect of canopy complexity within 4-species mixtures

The same situation as for generic diversity also seems to apply to comparisons of low vs high canopy complexity. There is seemingly no difference in AGB estimates, when holding species richness constant.

```
Rapideye_data %>%
    filter(Spp_richness == 4) %>%
    group_by(Canopy_type) %>%
    summarise(Mean_Biomass = mean(Biomass), Se_Biomass = std.error(Biomass)) %>%
    ggplot(aes(x = Canopy_type, y = Mean_Biomass)) + geom_pointrange(aes(ymin = Mean_Biomass -
    2 * Se_Biomass, ymax = Mean_Biomass + 2 * Se_Biomass,
    colour = Canopy_type), shape = Canopy_type), size = 1,
    fill = "white", shape = 22) + scale_colour_manual(values = cols_2) +
    theme(legend.position = "none") + labs(y = expression(atop("Aboveground biomass",
    paste("(Mg  ", Ha^-1, ")"))), x = "Canopy complexity") +
    scale_y_continuous(limits = c(225, 239), breaks = c(225,
        230, 235))
```

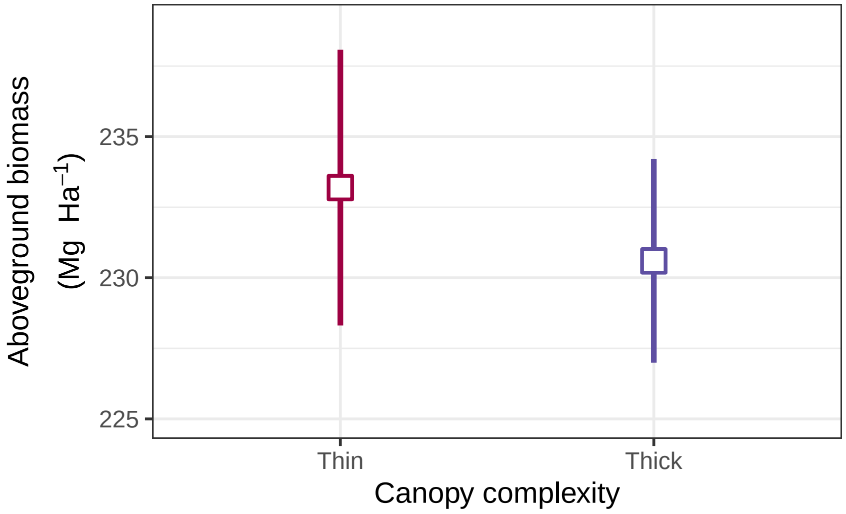

Figure 7: Aboveground biomass as a function of canopy complexity of plots planted with four-species (low vs high). Note: this figure relates to Fig. 1 (H) within the main manuscript.

##### 3.7 Effect of climber cutting

Within 16-species mixtures, there is no statistically detectable difference between plots which have and have not had a round of climber cutting as of 2012. The standard error associated with those which did undergo climber cutting is wider due to the smaller sample size compared to untreated 16-species mixtures (10 vs 38):

```
Rapideye_data %>%
    filter(Spp_richness == 16) %>%
    group_by(Climber_cutting) %>%
    summarise(Mean_Biomass = mean(Biomass), Se_Biomass = std.error(Biomass)) %>%
    ggplot(aes(x = Climber_cutting, y = Mean_Biomass)) +
    geom_pointrange(aes(ymin = Mean_Biomass - 2 * Se_Biomass,
        ymax = Mean_Biomass + 2 * Se_Biomass, colour = Climber_cutting,
        shape = Climber_cutting), size = 1, fill = "white",
        shape = 22) + scale_colour_manual(values = cols_2) +
    theme(legend.position = "none") + labs(y = expression(atop("Aboveground biomass",
    paste("(Mg  ", Ha^-1, ")"))), x = "Liana removal") +
    scale_y_continuous(limits = c(254, 273), breaks = c(256,
        260, 264, 268, 272))
```

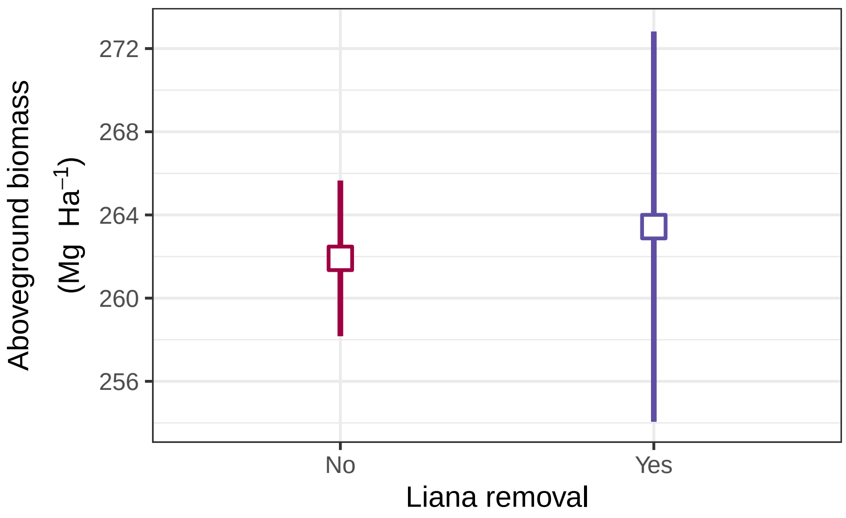

Figure 8: Aboveground biomass as a function of liana removal for plots with sixteen-species (climber cutting vs control). Note: this figure relates to Fig. 1 (G) within the main manuscript.

### 4 Statistical Modelling

Through the iterative model building process (Pinheiro & Bates 2002) we created two linear mixed effects models describing the growth of our plots via the RapidEye satellite indices. This approach attempts to reduce multiple testing using *a priori* contrasts within the overall model to test the more specific research questions:

#### 4.1 `me1_mass` model fitting

In `me1_mass`, we first look at the overall effect of all 5 treatments. We use *a priori* contrasts for replanting (replanted vs. not), the effect of species richness of the enrichment planted trees. We began by fitting `Planting` as our first fixed effect. This effectively splits the data into planted and unplanted groups. By subsequently fitting `Spp_richness` we then look at the effect of species richness (1, 4, or 16).

We also performed *a priori* contrasts for the treatments nested within the 4-species level. Specifically, we fit `Gen_div` and `Canopy_type` to look at the effects of the manipulations of these treatments within the 4-species planting treatment level.

Finally, we fit the `Treatment` fixed effect. All `Spp_richness` values have just one level except 16-species mixtures, which have `16_spp` and `16_cut`, allowing this fixed effect to exclusively explain variation within 16-species combinations. This specific ordering of all of our models in the sequential model allows contrasts for the effects of planting, species richness, generic diversity, canopy complexity, and liana removal (referred to as `Treatment` within our models) to be explored within a single initial overall model.

The mixed-effects model includes random effects for block [`(1|Block)`] and the different species compositions [`(1|Spp_comp)`].

```
me1_mass <- lmer(Biomass ~ Planting + factor(Spp_richness) +
    Gen_div + Canopy_type + Treatment + (1 | Block) + (1 |
    Spp_comp), data = Rapideye_data)
```

#### 4.2 `me2_mass` model fitting

`me2_mass` is a modification `me1_mass` which allows us to fit a linear regression ('linear contrast') for species richness, which is not possible in `me1_mass` due to the presence of unplanted controls having a `Spp_richness` value of zero, which prevents the $\log_2$ transformation. The initial effect of the species richness factor from model me1_mass is divided into a (log-) linear contrast (linear regression on $\log_2$ species richness) and the deviation from linearity (by fitting the species richness factor *after* the linear term).

As `me1_mass` provides values for planted vs unplanted controls, and all other fixed effects do not explore any further variation within unplanted controls, in `me2_mass` we subset the data to remove these values, which allows us to fit the regression on the $\log_2$ scale. All other fixed effects remain in the same order for the same reasons as explained above.

```
me2_mass <- lmer(Biomass ~ log2(Spp_richness) + factor(Spp_richness) +
    Gen_div + Canopy_type + Treatment + (1 | Block) + (1 |
    Spp_comp), subset = Spp_richness > 0, data = Rapideye_data)
```

Type I analysis of variance with Satterthwaite's method was applied to each model using the lmerTest package (version 3.1-3).

#### 4.3 Fit of model `me1_mass` and effect of planting

Within `me1_mass`, there is an effect of replanting with plots enrichment planted with a single species having 32.0 Mg ha$^{-1}$ greater AGB estimates than unplanted controls. Additionally, we see a highly significant effect of planted species richness influencing aboveground biomass.

```
anova(me1_mass, type = "I")
```

```
## Type I Analysis of Variance Table with Satterthwaite's method
##                      Sum Sq Mean Sq NumDF  DenDF  F value Pr(>F)
## Planting              35420   35420     1 116.00 222.6623 <2e-16
## factor(Spp_richness)  47983   23991     2 116.03 150.8178 <2e-16
## Gen_div                  89      89     1 116.00   0.5571 0.4569
## Canopy_type              54      54     1 116.00   0.3389 0.5616
## Treatment               128     128     1 116.29   0.8059 0.3712
```

```
display(me1_mass)
```

```
## lmer(formula = Biomass ~ Planting + factor(Spp_richness) + Gen_div +
##     Canopy_type + Treatment + (1 | Block) + (1 | Spp_comp), data = Rapideye_data)
##                       coef.est coef.se
## (Intercept)           182.67     4.43
## PlantingPlanted        83.06     5.51
## factor(Spp_richness)1 -51.10     7.86
## factor(Spp_richness)4 -33.47     5.65
## Gen_div2               -3.33     4.46
## Canopy_typeThin         2.60     4.46
## Treatment16_spp        -4.18     4.65
##
## Error terms:
##  Groups   Name        Std.Dev.
##  Spp_comp (Intercept)  0.00
```

```
##   Block    (Intercept)  3.58
##   Residual             12.61
## ---
## number of obs: 124, groups: Spp_comp, 35; Block, 2
## AIC = 966.4, DIC = 1006.6
## deviance = 976.5
```

##### 4.4 Residual diagnostics for `me1_mass`

The residual plots show no problematic patterns in residual variation. No transformation is required.

```
plot(me1_mass)
```

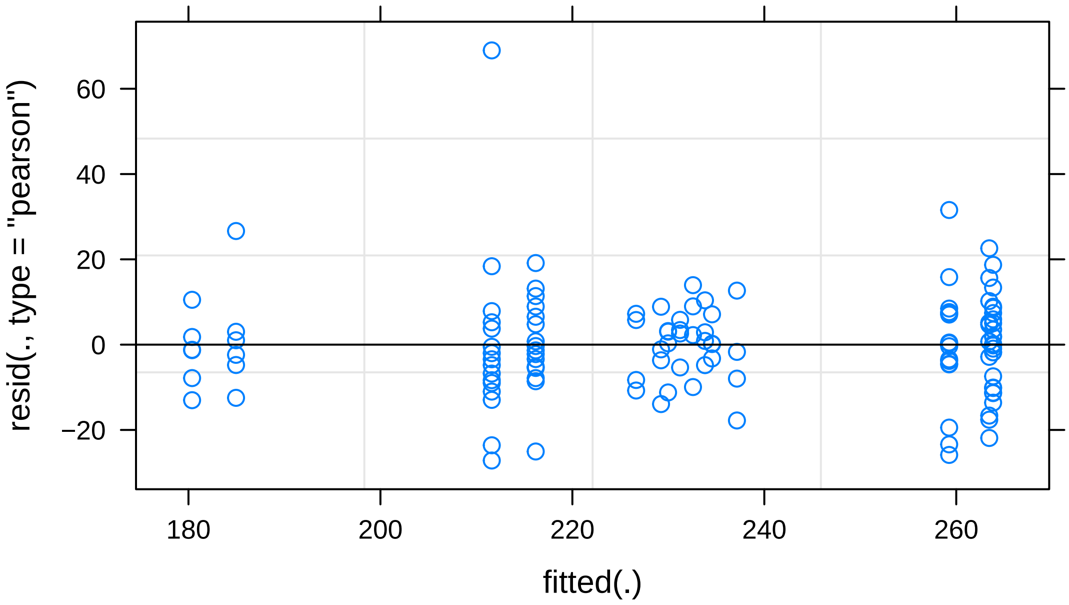

##### 4.5 Fit of model `me2_mass` and effect of species richness

Investigating this further, the majority of this variation can be explained through a regression with species richness expressed as a base 2 logarithm to better fit the assumptions of a linear mixed effects model. This model estimates that for every doubling in species richness a 13.54 Mg ha$^{-1}$ increase in AGB was observed after 10 years of establishment. Of the total variation explained by species richness, 98.1% is captured as the log2 regression curve, and the remaining 1.9% is deviation from this linearity.

Also, we see that regardless of the model we chose there was no observed effect of either generic diversity or canopy complexity on estimated aboveground biomass after accounting for species richness.

```
anova(me2_mass, type = "I")
```

```
## Type I Analysis of Variance Table with Satterthwaite's method
##                        Sum Sq Mean Sq NumDF  DenDF  F value  Pr(>F)
```

```
## log2(Spp_richness)    47057   47057     1 105.07 286.9738 < 2e-16
## factor(Spp_richness)    882     882     1 105.02   5.3778 0.02233
## Gen_div                  89      89     1 105.00   0.5405 0.46387
## Canopy_type              54      54     1 105.00   0.3287 0.56763
## Treatment               109     109     1 104.03   0.6665 0.41612
```

```
display(me2_mass)
```

```
## lmer(formula = Biomass ~ log2(Spp_richness) + factor(Spp_richness) +
##     Gen_div + Canopy_type + Treatment + (1 | Block) + (1 | Spp_comp),
##     data = Rapideye_data, subset = Spp_richness > 0)
##                       coef.est coef.se
## (Intercept)           213.89     3.23
## log2(Spp_richness)     13.54     1.64
## factor(Spp_richness)4  -9.44     3.99
## Gen_div4                3.33     4.53
## Canopy_typeThick       -2.60     4.53
## Treatment16_spp        -3.86     4.73
##
## Error terms:
##  Groups   Name        Std.Dev.
##  Spp_comp (Intercept)  0.00
##  Block    (Intercept)  3.26
##  Residual             12.81
## ---
## number of obs: 112, groups: Spp_comp, 34; Block, 2
## AIC = 880.6, DIC = 908.4
## deviance = 885.5
```

#### 4.6 Effect of generic diversity and canopy type

From within our four-species plots we did not identify any significant difference between plots with increased generic diversity. This is interesting as we have seen such a strong effect of species richness, however may be due to the questionable taxonomy of dipterocarps in Borneo (e.g. *Shorea* genus argued to be split into several smaller genera).

We also found no significant effect of canopy complexity on the observed biomass estimates. This again is surprising, but could be attributed to several causes, including that the experiment has not ran for long enough for the differences in canopy structure to develop (or satellite imagery is poor at estimating sub-canopy foliage and therefore biomass).

There was no statistically detectable impact of climber cutting observed between our plots in terms of biomass differences. This could again be attributed to several reasons. The most likely explanation is that at the time of the RapidEye data collection the liana removal treatment had only been applied once to one block (this treatment has subsequently been applied to both blocks and repeated).

#### 4.7 Residual diagnostics for `me2_mass`

The residual diagnostic plot for `me2_mass` is very similar to that of `me1_mass`, and shows no problematic patterns.

```
plot(me2_mass)
```

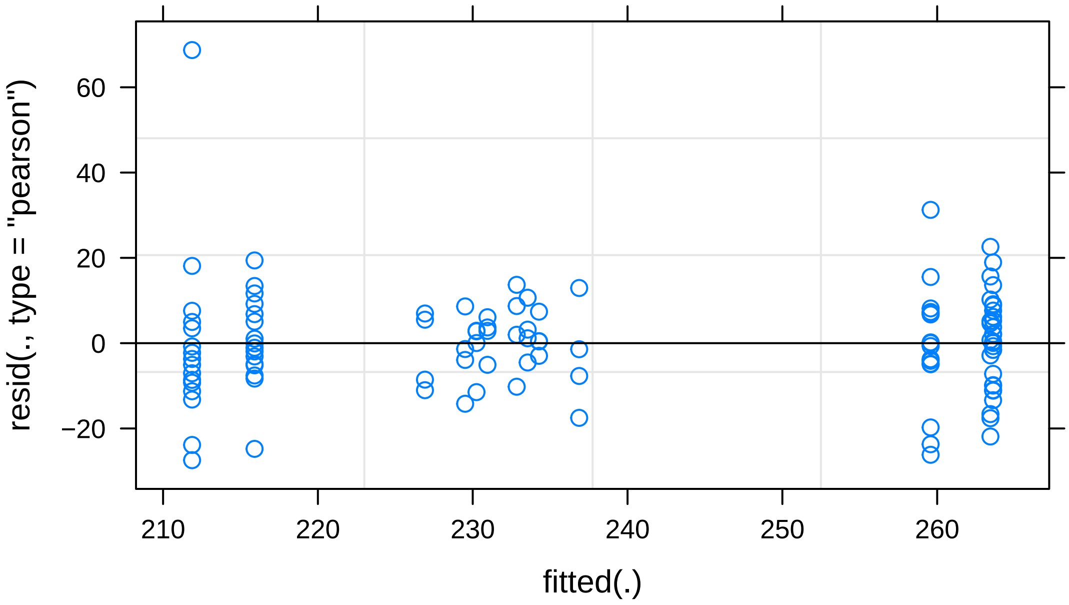

##### 4.8 me1_landsat model fitting

To account for how these effects change over time we separately analysed cover from our Landsat data using the following model which is based on the previous models but extended to include the three time points at which the Landsat data was collected.

```
me1_landsat <- lmer(Cover ~ Planting + factor(Year) * factor(Spp_richness) +
    Gen_div + Canopy_type + Treatment + (1 | Block) + (1 |
    Spp_comp) + (1 | Plot), data = Landsat_data)
```

##### 4.9 Fit of model me1_landsat

This model shows that there is a strong effect of both year and species richness, as well as an interaction between the two. This confirms the visual conclusions drawn earlier on.

```
anova(me1_landsat, type = "I")
```

```
## Type I Analysis of Variance Table with Satterthwaite's method
##                               Sum Sq Mean Sq NumDF DenDF   F value
## Planting                       33.53   33.53     1   117   62.0703
## factor(Year)                 1217.83  608.91     2   240 1127.0484
## factor(Spp_richness)           14.29    7.14     2   117   13.2227
## Gen_div                         0.68    0.68     1   117    1.2565
## Canopy_type                     0.10    0.10     1   117    0.1801
## Treatment                       1.75    1.75     1   117    3.2373
```

```
## factor(Year):factor(Spp_richness)  221.55   36.93     6   240   68.3464
##                                     Pr(>F)
## Planting                          1.894e-12
## factor(Year)                      < 2.2e-16
## factor(Spp_richness)              6.648e-06
## Gen_div                             0.26461
## Canopy_type                         0.67209
## Treatment                           0.07456
## factor(Year):factor(Spp_richness) < 2.2e-16
```

```
display(me1_landsat)
```

```
## lmer(formula = Cover ~ Planting + factor(Year) * factor(Spp_richness) +
##     Gen_div + Canopy_type + Treatment + (1 | Block) + (1 | Spp_comp) +
##     (1 | Plot), data = Landsat_data)
##                                       coef.est coef.se
## (Intercept)                           71.95     0.38
## PlantingPlanted                        1.03     0.54
## factor(Year)2005                       0.51     0.30
## factor(Year)2010                       0.56     0.30
## factor(Spp_richness)1                 -0.76     0.74
## factor(Spp_richness)4                 -0.66     0.54
## Gen_div2                               0.47     0.42
## Canopy_typeThin                       -0.18     0.42
## Treatment16_spp                       -0.76     0.42
## factor(Year)2005:factor(Spp_richness)1   2.42     0.35
## factor(Year)2010:factor(Spp_richness)1   2.13     0.35
## factor(Year)2005:factor(Spp_richness)4   2.42     0.35
## factor(Year)2010:factor(Spp_richness)4   4.00     0.35
## factor(Year)2005:factor(Spp_richness)16  3.45     0.34
## factor(Year)2010:factor(Spp_richness)16  5.57     0.34
##
## Error terms:
##  Groups   Name        Std.Dev.
##  Plot     (Intercept) 1.10
##  Spp_comp (Intercept) 0.00
##  Block    (Intercept) 0.00
##  Residual             0.74
## ---
## number of obs: 372, groups: Plot, 124; Spp_comp, 35; Block, 2
## AIC = 1120.2, DIC = 1048
## deviance = 1065.1
```

#### 4.10 Residual diagnostics for `me1_landsat`

The residual diagnostic plots show no problematic patterns in residual variation.

```
plot(me1_landsat)
```

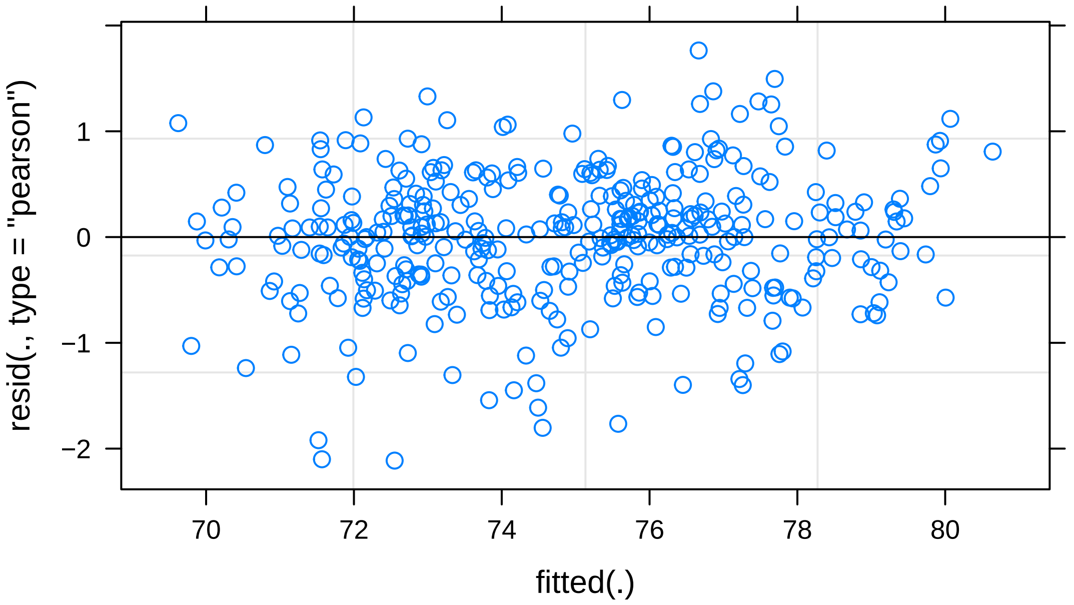

### 5 Phylogenetic and functional diversity measures

The effects, or lack thereof, of generic diversity on estimated aboveground biomass values led us to further investigate. To better understand why this may be we calculated a measure of phonetically diversity, Faith's PD, and functional diversity.

Loading in the following datasets:

- `comp_borneo` - community composition for traits at Borneo
- `pruned_tree` - the phylogeny tree used
- `all_spp_traits` - all available functional trait data on our dipterocarps
- `huang_traits` - trait data matching the traits referred to in Huang et al (2018)
- `communities` - unweighted version of `comp_Borneo`, which does not contain formation on planted species densities

```
comp_borneo <- readRDS("Data/borneo_plot_compositions.rds")
pruned_tree <- readRDS("Data/phylo_tree.rds")
all_spp_traits <- readRDS("Data/trait_info_borneo.rds")
huang_traits <- readRDS("Data/trait_info_huang_2018.rds")
communities <- readRDS("Data/borneo_plot_compositions_unweighted.rds")
```

#### 5.1 Phylogenetic diversity measure

Faith's phylogenetic diversity (PD) is a measurement dependent on the species richness of each plot. It uses a constructed phylogenetic tree to calculate it's total length. As such more species-rich trees have by default a greater score. This measurement is particularly good, therefore, for measuring phylogenetic diversity within the species level.

Calculating Faith's PD was done using the `pd` function in the `picante` package (version 1.8.2).

```
faith_pd_result <- pd(communities, pruned_tree, include.root = TRUE) %>%
    rownames_to_column(var = "Spp_comp") %>%
    select(-SR) %>%
    mutate(Spp_comp = as.factor(Spp_comp))

head(faith_pd_result)
```

```
##                      Spp_comp         PD
## 1 1_Dipterocarpus_conformis 0.01760607
## 2 1_Dryobalanops_lanceolate 0.01865573
## 3         1_Hopea_ferruginea 0.01663012
## 4             1_Hopea_sangal 0.01935996
## 5   1_Parashorea_tomentalla 0.01546887
## 6   1_Parashorea_malaanonan 0.01467378
```

#### 5.2 Graphical representation of phylogenetic diversity data

Graphing the relationship between Faith's PD and aboveground biomass scores we see that initially there is a positive relationship between the two, but this is lost when we focus specifically on the four-species plots.

```
data2 <- Sat_data %>%
    filter(Index == "Biomass" & Satellite == "RapidEye",
        Spp_richness == 4)
data3 <- Sat_data %>%
    filter(Index == "Biomass" & Satellite == "RapidEye")

Sat_data %>%
    filter(Index == "Biomass") %>%
    filter(Spp_richness != 4) %>%
    ggplot(aes(PD, Score)) + geom_point() + geom_point(data = data2,
    aes(colour = Gen_div, shape = Canopy_type)) + geom_smooth(data = data3,
    mapping = aes(x = PD, y = Score), method = "lm", se = FALSE) +
    geom_smooth(data = data2, mapping = aes(x = PD, y = Score),
        method = "lm", se = FALSE, linetype = 2, colour = "black") +
    labs(x = "Phylogenetic diversity (Faith's PD)", y = expression(paste("Aboveground biomass",
        " ", "(Mg h", a^-1, sep = "", ")")), shape = "Canopy\ntype",
        colour = "Generic\ndiversity") + scale_colour_manual(values = cols) +
    scale_shape_manual(values = c(15, 17)) + theme_bw()
```

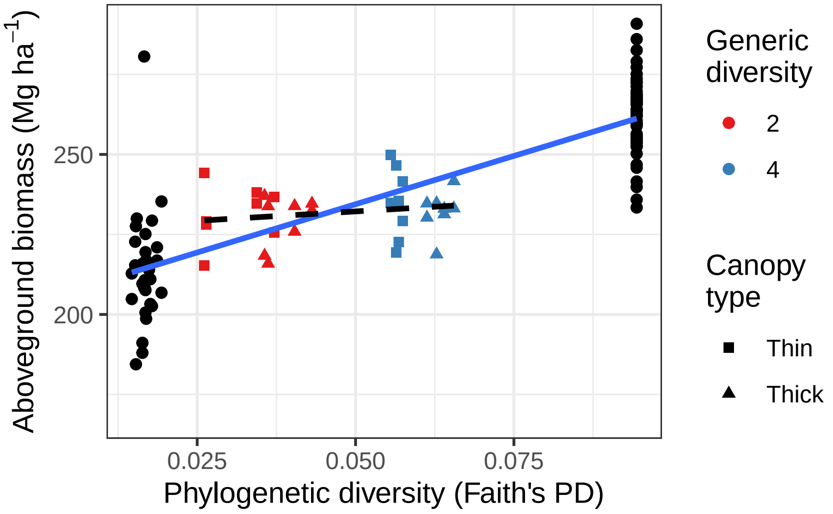

Figure 9: Measures of functional diversity increase across the full diversity gradient from 1 to 16 species but not in relation to the 5 treatments applied to the subset of four - species plots that manipulate generic diversity (2 vs. 4 genera) and canopy complexity (lower vs. higher). Note: This figure relates to Fig. 3 (B) within the main manuscript.

#### 5.3 me1_PD model fitting

Once species richness is accounted for, there is no effect of phylogenetic diversity to be seen within the 4-species groups.

```
me1_PD <- lmer(Biomass ~ log2(Spp_richness) + factor(Spp_richness) +
    PD + Canopy_type + Treatment + (1 | Block) + (1 | Spp_comp),
    subset = Spp_richness > 0, data = Rapideye_data)
```

```
anova(me1_PD, type = "I")
```

```
## Type I Analysis of Variance Table with Satterthwaite's method
##                      Sum Sq Mean Sq NumDF  DenDF  F value  Pr(>F)
## log2(Spp_richness)    47057   47057     1 105.07 287.9543 < 2e-16
## factor(Spp_richness)    882     882     1 105.02   5.3963 0.02211
## PD                       94      94     1 105.00   0.5726 0.45094
## Canopy_type             107     107     1 105.00   0.6577 0.41920
## Treatment               109     109     1 104.05   0.6696 0.41506
```

```
display(me1_PD)
```

```
## lmer(formula = Biomass ~ log2(Spp_richness) + factor(Spp_richness) +
##     PD + Canopy_type + Treatment + (1 | Block) + (1 | Spp_comp),
##     data = Rapideye_data, subset = Spp_richness > 0)
##                       coef.est coef.se
## (Intercept)           211.11     4.36
## log2(Spp_richness)     10.63     3.48
## factor(Spp_richness)4  -6.43     3.58
## PD                    165.74   174.66
```

```
## Canopy_typeThick       -3.81     4.70
## Treatment16_spp        -3.86     4.72
##
## Error terms:
##  Groups   Name        Std.Dev.
##  Spp_comp (Intercept)  0.00
##  Block    (Intercept)  3.26
##  Residual             12.78
## ---
## number of obs: 112, groups: Spp_comp, 34; Block, 2
## AIC = 873, DIC = 915.3
## deviance = 885.2
```

##### 5.4 Residual diagnostics for `me1_PD`

Besides a single large outlier this distribution of residuals seems reasonable again. It has the same limitations for other RapidEye-based models, but is still acceptable.

```
plot(me1_PD)
```

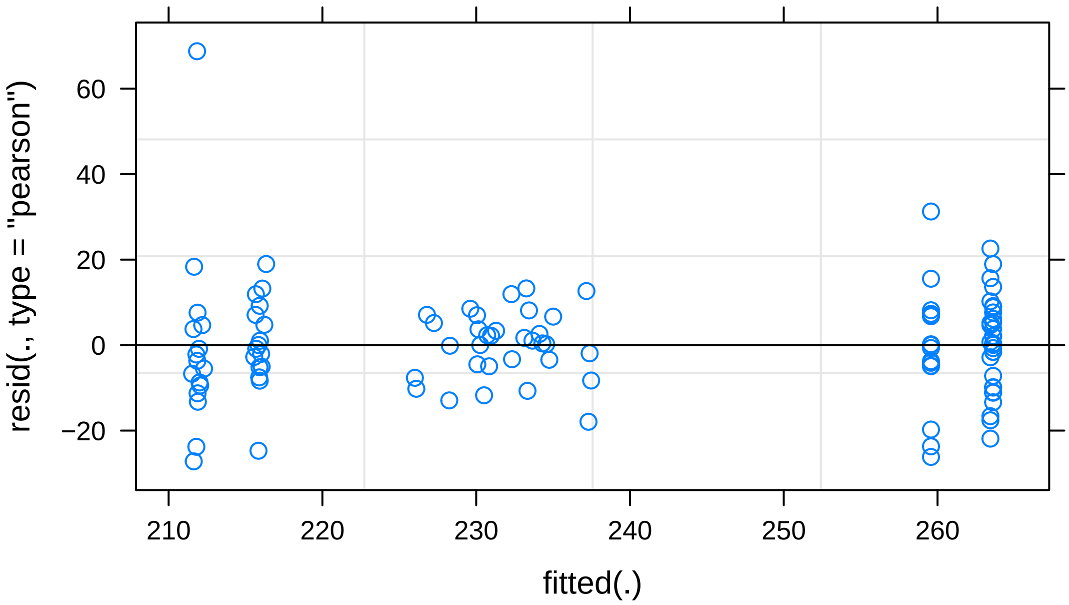

##### 5.5 Functional diversity measure

As there is evidence that the phylogeny of the Dipterocarpaceae is not yet fully accurate, casting doubt onto the effectiveness of using PD as a measure of diversity, we investigated if there was a relation between functional diversity and the aboveground biomass differences seen. These measurements may act as a reasonably good proxy for genetic diversity, whilst avoiding the errors of the current phylogeny.

Functional diversity is analogous to Faith's PD, in that it is dependent on species richness. This is as it sums up the total branch length of the 'functional dendrogram' we create. Following the methodology of Petchy & Gaston (2002) we use the following code to generate FD data:

```r
FD_function <- function(data) {

    comp_borneo_table <- comp_borneo %>%
        as.data.frame() %>%
        rownames_to_column("Spp_comp")

    names <- rownames(comp_borneo)  # extracting names of each species composition level

    table <- tibble(Spp_comp = "NA", species = "NA")

    # This loop creates a list of all species for
    # every given species composition
    for (i in 1:length(names)) {
        comp <- names[i]
        spp <- comp_borneo_table[i, ] %>%
            .[, colSums(. != 0) > 0] %>%
            select(-Spp_comp) %>%
            colnames()
        table2 <- tibble(Spp_comp = comp, species = toString(spp))
        table <- table %>%
            rbind(table2)

    }

    # Removing all monocultures as well as NA values
    # (so control removed too)
    table <- table %>%
        filter(species != "NA") %>%
        filter(!str_detect(Spp_comp, "1_"))

    # For each row in the table it will then
    # calculate the Gower distance based on the
    # species provided

    result <- tibble(Spp_comp = "NA", FD = "NA")

    for (i in 1:nrow(table)) {

        test_traits <- data[str_detect(table[[i, 2]], row.names(data)),
            ]

        a <- SumBL(test_traits, gower.dist = TRUE, method.hclust = "average",
            scale.tr = TRUE, method.dist = "euclidian")

        result2 <- tibble(Spp_comp = table[[i, 1]], FD = a)

        result <- result %>%
            rbind(result2) %>%
            filter(Spp_comp != "NA")

    }

    result
```

```
}
```

Although we have trait information for a much wider array of traits, calculating functional dispersion for all available traits results in poor distinction between species.

```
FD_function(all_spp_traits)
```

```
## # A tibble: 17 x 2
##    Spp_comp FD
##    <chr>    <chr>
##  1 4_1      2
##  2 4_2      2
##  3 4_3      2.11620217126183
##  4 4_4      2.16610186815281
##  5 4_5      2
##  6 4_6      2
##  7 4_7      2
##  8 4_8      2.07492606034256
##  9 4_9      2
## 10 4_10     2.08225852367672
## 11 4_11     2.12041854789981
## 12 4_12     2.08416321389922
## 13 4_13     2.14020543087222
## 14 4_14     1.961149614756
## 15 4_15     2.10484173842503
## 16 4_16     1.98034247975787
## 17 16_spp   3.53750052726983
```

The traits we will use are those which have a demonstrated impact on the growth/survival trade off seen in dipterocarps. They are also the traits used in Huang et al. (2018), a related paper. As such we continue the analysis with these traits (referred to as `huang_traits` for convenience).

```
FD_result_huang <- FD_function(huang_traits)
FD_result_huang
```

```
## # A tibble: 17 x 2
##    Spp_comp FD
##    <chr>    <chr>
##  1 4_1      2.16542623853604
##  2 4_2      2.14319436308913
##  3 4_3      2.03327335284352
##  4 4_4      2.05312045517076
##  5 4_5      2.09179389706832
##  6 4_6      2.04504410785619
##  7 4_7      2.07488559748087
##  8 4_8      2.09737457359865
##  9 4_9      2.19180395797017
## 10 4_10     2.13775647860784
## 11 4_11     2.17389730769747
## 12 4_12     2.07888675678027
## 13 4_13     2.1950965389224
```

```
## 14 4_14     1.99226840057852
## 15 4_15     2.04498841143234
## 16 4_16     2.04235968749777
## 17 16_spp   3.5935660347704
```

#### 5.6 Graphical representation of functional diversity data

The same trend as with phylogenetic measures is seen for functional trait measurements. Although a significant effect of functional diversity is seen for overall biomass, once we eliminate the effect of species richness this result is lost.

```
data2 <- Sat_data %>%
    filter(Index == "Biomass" & Satellite == "RapidEye",
        Spp_richness == 4)
data3 <- Sat_data %>%
    filter(Index == "Biomass" & Satellite == "RapidEye")

Sat_data %>%
    filter(Index == "Biomass") %>%
    filter(Spp_richness != 4) %>%
    ggplot(aes(FD, Score)) + geom_point() + geom_point(data = data2,
    aes(colour = Gen_div, shape = Canopy_type)) + geom_smooth(data = data3,
    mapping = aes(x = FD, y = Score), method = "lm", se = FALSE) +
    geom_smooth(data = data2, mapping = aes(x = FD, y = Score),
        method = "lm", se = FALSE, linetype = 2, colour = "black") +
    labs(x = "Functional diversity (FD)", y = expression(paste("Aboveground biomass",
        " ", "(Mg h", a^-1, sep = "", ")")), colour = "Generic\ndiversity",
        shape = "Canopy\ntype") + scale_colour_manual(values = cols) +
    scale_shape_manual(values = c(15, 17)) + theme_bw()
```

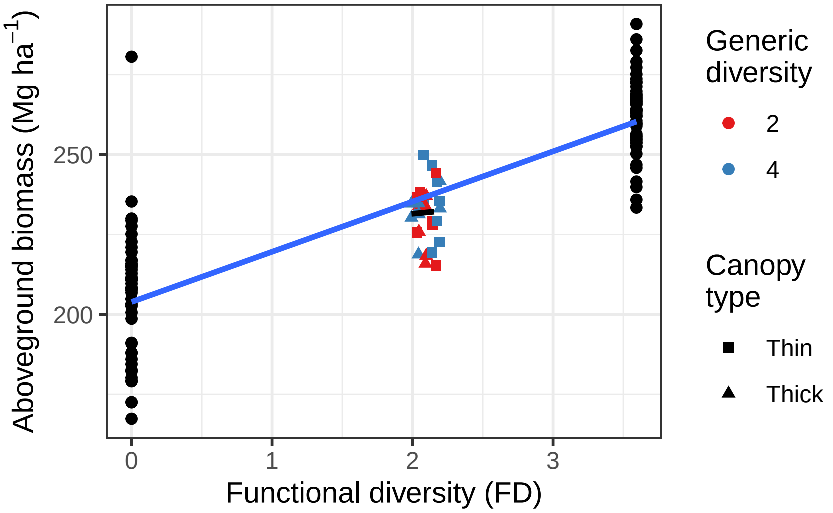

Figure 10: Measures of phylogenetic diversity (Faith's PD) increase across the full diversity gradient from 1 to 16 species but not in relation to the 5 treatments applied to the subset of four - species plots that manipulate generic diversity (2 vs. 4 genera) and canopy complexity ( lower vs. higher ). Note: This figure relates to Fig. 3 (A) within the main manuscript.

#### 5.7 me1_FD model fitting

Similarly to measures of PD, there is no discernible effect of FD after species richness has been accounted for.

```
me1_FD <- lmer(Biomass ~ log2(Spp_richness) + factor(Spp_richness) +
    FD + Canopy_type + Treatment + (1 | Block) + (1 | Spp_comp),
    subset = Spp_richness > 0, data = Rapideye_data)
```

```
anova(me1_FD, type = "I")
```

```
## Type I Analysis of Variance Table with Satterthwaite's method
##                      Sum Sq Mean Sq NumDF  DenDF  F value  Pr(>F)
## log2(Spp_richness)    47056   47056     1 105.07 285.5470 < 2e-16
## factor(Spp_richness)    882     882     1 105.02   5.3509 0.02266
## FD                        2       2     1 105.00   0.0128 0.91026
## Canopy_type              55      55     1 105.00   0.3311 0.56626
## Treatment               109     109     1 104.00   0.6621 0.41768
```

```
display(me1_FD)
```

```
## lmer(formula = Biomass ~ log2(Spp_richness) + factor(Spp_richness) +
##     FD + Canopy_type + Treatment + (1 | Block) + (1 | Spp_comp),
##     data = Rapideye_data, subset = Spp_richness > 0)
##                         coef.est coef.se
## (Intercept)             213.89     3.23
## log2(Spp_richness)       18.35    37.19
## factor(Spp_richness)4    -6.19    12.71
## FD                       -5.28    40.80
## Canopy_typeThick         -2.86     4.96
## Treatment16_spp          -3.86     4.74
##
## Error terms:
##  Groups   Name        Std.Dev.
##  Spp_comp (Intercept)  0.00
##  Block    (Intercept)  3.26
##  Residual             12.84
## ---
## number of obs: 112, groups: Spp_comp, 34; Block, 2
## AIC = 876.8, DIC = 913.4
## deviance = 886.1
```

#### 5.8 Residual diagnostics for me1_FD

This is very similar to the phylogenetic model's residuals. Again, no need for changes to be made based on the results of this plot.

```
plot(me1_FD)
```

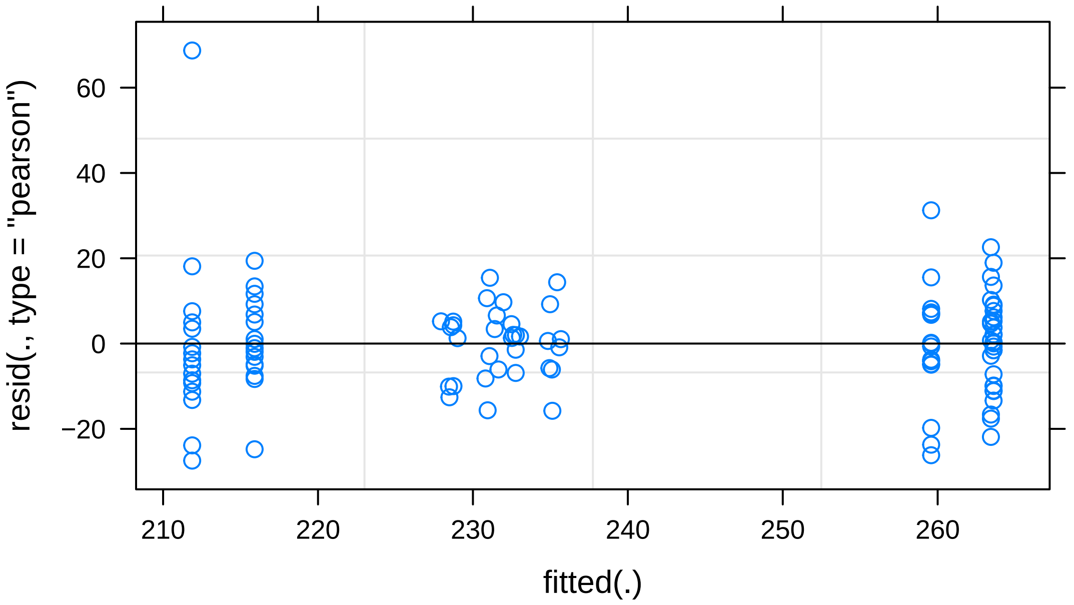
resid(., type = "pearson")
fitted(.)
60
40
20
0
−20
210
220
230
240
250
260
